# Recapitulating Patient-to-Patient Colorectal Cancer Tumor Heterogeneity Using Patient-Derived Xenograft Cells in an Engineered Tissue Model

**DOI:** 10.64898/2025.12.15.693817

**Authors:** Iman Hassani, Benjamin Anbiah, Yuan Tian, Bulbul Ahmed, William J. Van Der Pol, Elliot J. Lefkowitz, Peyton Kuhlers, Nicole L. Habbit, Martin J. Heslin, Michael W. Greene, Elizabeth A. Lipke

## Abstract

Establishing *in vitro* cancer models that more closely recapitulate patient tumor microenvironmental heterogeneity, including variations in stromal cells and mechanical properties that influence colorectal cancer (CRC) progression, is crucial for advancing CRC research. This study evaluated the ability of 3D engineered CRC-PDX (3D-eCRC-PDX) tissues to recapitulate the heterogeneity found between patient-derived xenograft (PDX) tumors from three CRC patients (stage II, III-B, and IV). To form the 3D-eCRC-PDX tissues, CRC-PDX tumor cells were encapsulated in PEG-fibrinogen hydrogels and maintained for 29 days *in vitro*. 3D-eCRC-PDX tissues recapitulated key patient-specific tumor characteristics. During long-term culture, 3D-eCRC-PDX tissues mimicked the patient-specific growth rates of the originating CRC-PDX tumors. Importantly, tumor cellular subpopulations, including the ratio of human cancer cells to mouse stromal cells and the ratios of proliferative human cancer cells and CK20^+^ cells were maintained in 3D-eCRC-PDX tissues, unlike in 2D cell culture. Differences in mechanical stiffness between the originating CRC-PDX tumors were also recapitulated by the 3D-eCRC-PDX tissues. Principal component analysis of transcriptomic data clustered 3D-eCRC-PDX tissues and CRC-PDX tumors together by patient, indicating similar gene expression profiles. These findings highlight the potential of 3D-eCRC-PDX tissues as a promising tool for CRC research, capable of maintaining patient-specific tumor microenvironment heterogeneity.

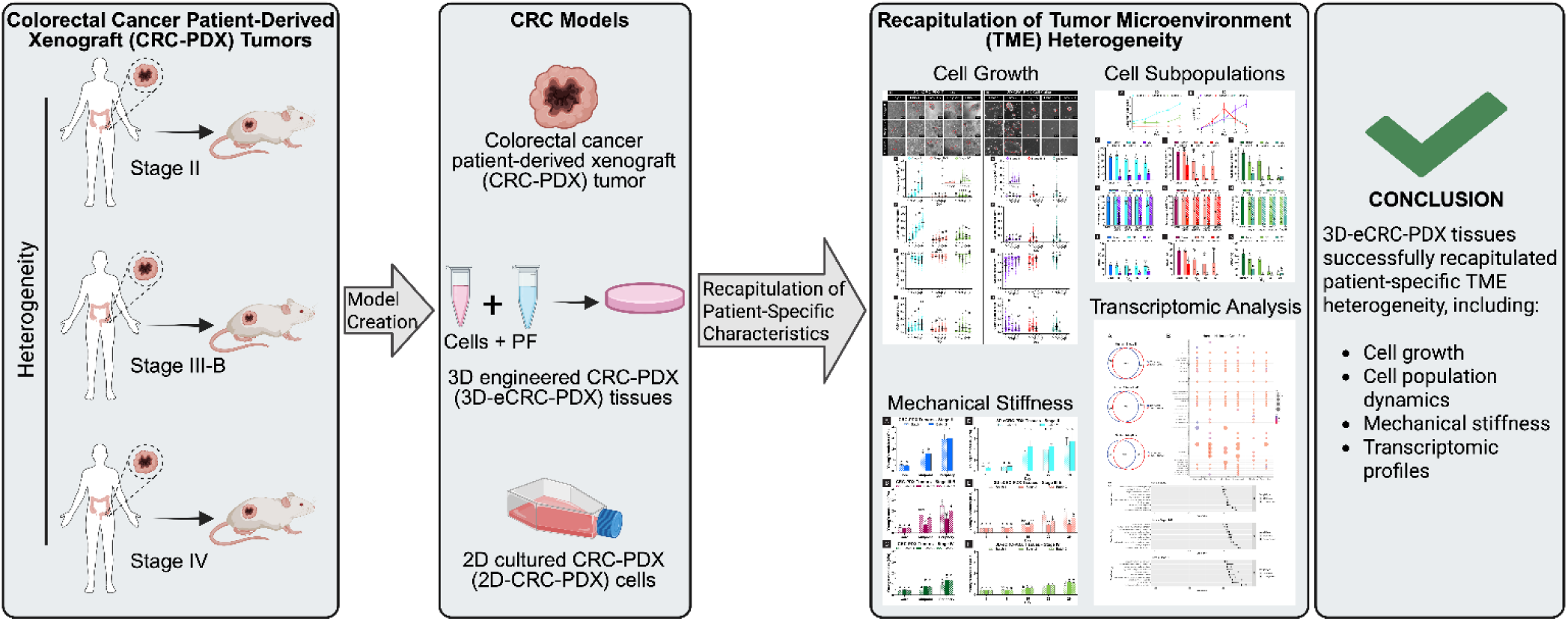

**Statement of Significance:** This study establishes engineered colorectal cancer tissues formed using PEG-fibrinogen and patient-derived xenograft (PDX) tumor cells for modeling inter-patient tumor heterogeneity. In vitro models that preserve patient tumors’ biological and structural heterogeneity are central to the development of more predictive and reproducible preclinical cancer models. Here engineered tissues replicated patient-specific tumor growth dynamics, sustained key cancer and stromal cell subpopulations, recapitulated originating PDX tumor stiffness, and sustained patient-specific patterns of gene expression. This work demonstrates the long-term culture of cells from patient xenografts in engineered colorectal cancer tissues, enabling sustained preservation of critical patient tumor-specific microenvironmental characteristics in vitro.

## Introduction

In the United States, colorectal cancer (CRC) ranks as the fourth most frequently diagnosed malignancy and is responsible for the second-highest number of cancer-related deaths [1, 2]. To understand CRC biology and address relevant research questions, development of cancer models that better recapitulate tumor heterogeneity is needed. Although significant insights into CRC have been achieved through the use of modeling, the majority of the current *in vitro* models have focused on the use of standard cell lines [3–10]. As cell lines are typically limited to one cell subpopulation and selectively isolated during establishment, they lack the extensive heterogeneity observed in patients’ tumors [11, 12]. To overcome this deficiency, patient-derived cells can be grown in immunocompromised mice [13] in what is described as a patient-derived xenograft (PDX).

PDX models more accurately recapitulate the heterogeneity of patients’ tumors as compared to standard cell lines [11, 14]. Evidence indicates that PDX tumors grown in immunocompromised mice generally retain the molecular features, transcriptomic signatures, and genetic alterations observed in the corresponding patient tumors [15–17]. Although these PDX models are a useful tool to examine tumor biology, the use of PDX models involves extensive animal handling, thereby resulting in a costly, time-consuming, and low throughput platform [18–20]. To address these limitations, several research groups have attempted to culture PDX cells in *in vitro* 2D models [21, 22]. However, this transition of PDX cells to a 2D monolayer culture alters the balance of cell types present, ultimately yielding an *in vitro* cancer model that fails to replicate the complex subpopulations found *in vivo* [21, 22]. Other groups utilized patient-derived cells in *in vitro* 3D CRC spheroid or organoid models [23–28]. Although these studies have advanced model development, the resulting systems tend to maintain only a fraction of the cellular subtypes present in the PDX tumor. The inclusion of multiple subpopulations, especially stromal cells [29], better mirrors the complex heterogeneity of the parental tissue. In addition, organoid and self-aggregated spheroid models provide limited capacity to recapitulate or modulate the TME stiffness, which has been correlated with disease progression and survival and, therefore, considered a key aspect of modeling the TME.

Tissue engineering of CRC has the capacity to support tumor heterogeneity and more closely mimic patient tumors as compared to traditional 2D models by providing a biomechanical architecture that better represents the native tumor microenvironment [23–25, 30–39]. Biomaterial-based 3D models with defined compositions have been developed for colorectal cancer (CRC) modeling. Natural polymer systems such as hyaluronan–gelatin hydrogels have been shown to support CRC organoid–fibroblast co-culture [40], while chitosan-based cryogels have been used to demonstrate that scaffold stiffness regulates tumor morphology [41]. In our prior work, we formed high density (initially 20 million cells/cm^3^) engineered CRC tissues using the natural-synthetic hybrid material poly(ethylene glycol)-fibrinogen (PEG-Fb) and demonstrated recapitulation of CRC cell line-dependent phenotypic differences during long-term culture [38]. Additional approaches have also been established including decellularized ECM to preserve native biochemical cues [42, 43], 3D bioprinting to enable fabrication of composite hydrogels that modulate cell signaling [44], microfluidic tumor-on-a-chip systems that incorporate matrices [45, 46], and heterotypic co-culture platforms that incorporate stromal cells to reflect tumor microenvironment complexity [47]. However, most biomaterial-based 3D CRC models utilize established cell lines rather than patient-derived cells, limiting their ability to recapitulate patient-to-patient tumor heterogeneity. Additionally, many systems focus on short-term culture periods (7-14 days), which are suitable for drug screening but insufficient for studying long-term tumor behavior.

To more accurately replicate the patient tumor microenvironment, patient-derived cells and biomimetic materials such as hydrogels have been combined to create *in vitro* 3D tissue-engineered models for several types of cancers, including prostate [33, 48], breast [49, 50], and glioblastoma tissues [34–36]. In these studies, the synergic use of 3D scaffolds and patient-derived cells has enhanced the recapitulation of the native tumor microenvironmental cues, in some cases for longer *in vitro* PDX cell maintenance and propagation. In our previous work we established the ability to form engineered CRC tissues using PEG-Fb and CRC PDX tumor-derived cells and demonstrated proof-of-concept with a single CRC PDX line [51]. However, it remained uncertain whether tissue-engineered constructs could preserve the interpatient heterogeneity of tumors during extended in vitro culture, especially in the context of colorectal cancer (CRC).

To address this important question, in this study we utilized three PDX tumor lines derived from patients with stage II, III-B, and IV colorectal cancer (CRC) to generate engineered PDX cancer tissues by encapsulating PDX tumor cells within PEG-Fb hydrogels. Building on our prior work with one PDX line, here we evaluated how effectively this biomimetic platform captured the patient-to-patient heterogeneity of the original PDX tumors. To assess recapitulation of relative growth rates, changes in CRC PDX cell colony sizes and cell numbers in 3D engineered CRC PDX (3D-eCRC-PDX) tissues were quantified during long-term culture and compared to differences in the temporal changes in size of the originating CRC-PDX tumors. To evaluate maintenance of parental tumor cell compositions, cellular subpopulations in the 3D-eCRC-PDX tissues and 2D-CRC-PDX cell cultures were assessed throughout long-term (29 days) *in vitro* culture for all CRC-PDX lines. To examine the ability of the 3D-eCRC-PDX tissues to re-establish the differing biomechanical tumor microenvironments of the originating CRC-PDX tumors, line-dependent changes in the 3D-eCRC-PDX tissue mechanical stiffness were evaluated over time. To evaluate molecular fidelity, transcriptomic profiling with CMS classification of the 3D-eCRC-PDX tissues and CRC-PDX tumors was conducted. In addition to comparing the ability of the 3D-eCRC-PDX tissues and 2D-CRC-PDX cell cultures to recapitulate the differences between the originating parental CRC-PDX tumor lines, batch-to-batch reproducibility of 3D-eCRC-PDX tissues formed from each line was also assessed. Importantly, our results demonstrate that the 3D-eCRC-PDX tissues faithfully reproduce key features of the parental CRC-PDX tumors—including growth dynamics, cell subpopulation ratios, mechanical properties, and transcriptomic profiles—supporting their potential to model patient-to-patient tumor variability.

## Materials and Methods

### Preparation of PEG-Fb Hydrogels

Following established procedures [52, 53], poly(ethylene glycol) diacrylate (PEGDA) was synthesized, and fibrinogen was subsequently linked to PEG as described previously [54]. The PEG-Fb construct was characterized by examining both the efficiency of PEGylation and its ability to form 3D hydrogels through photocrosslinking [55].

### In Vivo Expansion of CRC PDX Cells

All animal work was conducted under the approval of the Auburn University Animal Care and Use Committee. Stage II, III-B, and IV CRC adenocarcinoma PDX lines, obtained frozen from BioConversant, LLC, were used to establish xenografts in NOD-SCID mice (The Jackson Laboratory) as previously described [37]. For the experiments described here, CRC-PDX tumors were propagated without Matrigel and passaged three to four times. In compliance with the IACUC protocol, tumors were not allowed to exceed 17 mm² (approximately 2.5 g if extrapolated to 17 mm³), and no tumors surpassed this limit.

### Preparation of CRC-PDX Cells for 2D and 3D Culture Systems

Tumors from stage II, III-B, and IV CRC-PDX models were harvested and dissociated following published methods [37]. The resulting PDX cells were isolated from each line and subsequently used to establish 2D cultures and construct 3D-eCRC-PDX tissues.

### Generation of 3D-eCRC-PDX Models

CRC-PDX cells isolated from each tumor line were encapsulated in PEG-Fb hydrogels at a density of 20 million cells/mL to form the 3D-eCRC-PDX tissues according to the previously established methods [37]. These constructs were cultured for 29 days at 37 °C in 5% CO₂, with medium refreshed every 2–3 days. Culture medium consisted of DMEM (Lonza) supplemented with 10% (v/v) FBS (Atlanta Biologicals™), 1% (v/v) pen/strep solution (HyClone™), and 2 mM glutaGRO™ Supplement (Corning).

### Culture and Growth Analysis of 2D CRC-PDX Cells

Isolated CRC cells from each CRC-PDX tumor line were plated in T-75 flasks and maintained under standard culture conditions (37 °C, 5% CO₂) as described previously [37]. Media were refreshed every 2–3 days. An initial passage was performed on Day 8 before cells reached complete growth arrest, based on: (1) visual assessment of cell confluence and (2) a standardized interval of a 7-day timepoint for comparison with 3D cultures. Passaging was performed using 0.25% trypsin-EDTA for 5 minutes at 37°C, with 6×10^5^ cells per T75 cell culture flask. Subsequent passages were performed every 7 days for the remainder of the 29-day culture period and analyses were conducted as described below.

To quantitatively assess growth kinetics without cell passaging, parallel cultures were established by seeding cells into ten T-25 flasks for each tumor line. One flask was harvested every 24 hours, enabling daily enumeration of cell number from Day 1 through Day 11.

### Assessment of Cell Viability and Imaging Workflow

Live/dead assay, confocal imaging, image processing, and viability quantification were performed according to the procedures as previously described [37].

### Morphological Assessment of CRC-PDX Cells in 3D and 2D Models

The 3D-eCRC-PDX tissues from each CRC-PDX tumor line were maintained *in vitro* for 29 days, with imaging performed every 7 days. Phase-contrast and Z-stack images were used to measure colony morphology, including area, axes, circularity, and aspect ratio, according to established methods [37]. Phase-contrast images of 2D-CRC-PDX cells were collected before each passage.

### Flow Cytometric Analysis of CRC-PDX Cell Subpopulations

Single-cell suspensions were prepared from CRC-PDX tumors, 3D-eCRC-PDX tissues, and 2D-CRC-PDX cultures to analyze cell subpopulations by flow cytometry. Viable cells were manually counted after dissociation. Human- and mouse-derived cells, CRC cells, and proliferating cells were identified using the following antibodies: B2M (mouse IgG2a mAb, OriGene Technologies) conjugated to Zenon™ PE, H-2 Db (mouse IgG2a mAb, OriGene Technologies) conjugated to Zenon™ PE, CK20 (rabbit IgG mAb, Cell Signaling Technology) conjugated to Zenon™ Alexa Fluor™ 647, and Ki-67 (rabbit IgG pAb, Abcam) conjugated to Zenon™ Alexa Fluor™ 647. All staining procedures including unstained cells, isotype, and dead cells controls and cell subpopulation quantifications were performed as previously described [37].

### Cell Morphology Assessment via Staining

To investigate cellular morphology in 3D-eCRC-PDX tissues, cells were stained for nuclei, CRC-specific markers, and F-actin filaments using Hoechst 33342, CK20-Alexa Fluor™ 488, and Alexa Fluor™ 568 Phalloidin (ThermoFisher Scientific), respectively. The staining and imaging procedures were performed as previously described in detail [37].

### Whole-Tissue Morphological Assessment of 3D-eCRC-PDX Constructs

To monitor temporal morphological changes in 3D-eCRC-PDX tissues, images were acquired from two perspectives: the side view using a CellScale MicroTester (previously called “MicroSquisher”) and the top view via phase-contrast microscopy. ImageJ software was employed to calculate tissue volume (both views), mean geometric diameter (top view, Equation 1), circularity (side view, Equation 2), and aspect ratio (side view, Equation 3). Analyses included at least three samples per time point per batch.

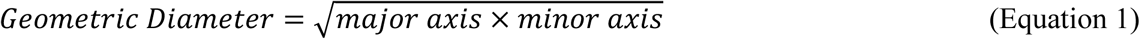

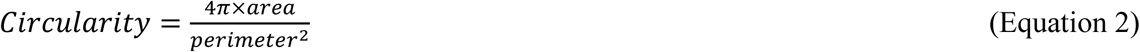

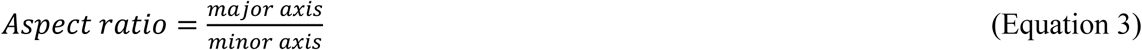

### Stiffness Analysis of CRC-PDX Tumors and Engineered 3D Tissues

The mechanical properties of CRC-PDX tumors and 3D-eCRC-PDX tissues were assessed using a CellScale MicroTester system in combination with SquisherJoy software [37]. For 3D-eCRC-PDX tissues, stiffness was measured at Days 1, 8, 15, 22, and 29. Each condition included at least three samples per batch.

### Immunostaining of Sectioned 3D-eCRC-PDX Tissues

3D-eCRC-PDX tissues (Days 15 and 22) were embedded in Tissue-Tek O.C.T. Compound (Electron Microscopy Sciences, Hatfield, PA), flash frozen, and stored at −80°C. Subsequently, tissue blocks were sectioned using a microtome cryostat (Microm HM 505E) at a thickness of 50 μm; this thickness was selected to preserve 3D tissue architecture for immunostaining analyses. To begin the immunostaining protocol, excess surrounding O.C.T. compound was gently rinsed away using PBS at room temperature. A fixation solution (4% paraformaldehyde (Electron Microscopy Sciences, Hatfield, PA) in PBS) was then added to each tissue section and incubated for one hour at room temperature.

After rinsing with PBS, tissue sections were stained with wheat germ agglutinin (WGA) at a 1:200 dilution for one hour at room temperature to label cellular membrane glycoproteins. After rinsing with PBS, tissue section cells were permeabilized with PBS-T (sterile filtered PBS with 0.2% v/v Triton X-100 and 1% FBS) for 15 minutes, then washed three times with PBS. Next, tissue sections were incubated in blocking buffer (3% FBS in PBS) overnight at 4°C. The following day, tissue sections were labeled with the MAB1273, Clone 113-1 antibody (Millipore Sigma, Burlington, MA) at a 1:200 dilution for three and a half hours at room temperature. Three PBS washes (10 minutes each) were performed, and the anti-Mouse IgG-Alexa Fluor 568 secondary antibody (Invitrogen, Waltham, MA) was added to the tissue sections at a 1:300 dilution for four hours at room temperature. Three, 10-minute PBS washes were again performed before counterstaining with the Hoechst 33342 compound (Millipore Sigma, Burlington, MA) at a 1:100 dilution for one hour at room temperature. Again, three PBS washes (10 minutes each) were performed. Serial ethanol drying was performed to prepare the stained tissue sections for mounting; 50% ethanol was added for five minutes, 75% ethanol was added for three minutes, 95% ethanol was added for two minutes, and 100% ethanol was added for two minutes followed by a two-minute air drying period. Stained tissue sections were mounted using the ProLong Gold Antifade Mountant (ThermoFisher Scientific, Waltham, MA) and a coverslip; clear nail polish was used to seal the coverslip edges following a 24-hour curing period. Finally, stained tissue sections were imaged using confocal microscopy (Nikon AI Confocal Scanning Laser Microscope) and the resultant fluorescent images were analyzed utilizing ImageJ software.

### Transcriptomic Analysis

Frozen CRC-PDX tumors representing Stage II (25–38 days), Stage III-B (42–125 days), and Stage IV (46–55 days), along with Day-15 3D-eCRC-PDX tissues, were processed for RNA extraction using RNeasy Plus Micro Kits (Qiagen). Quantification of total RNA was performed using a Qubit Fluorometer (Invitrogen), and RNA quality was evaluated using an Agilent 2100 Bioanalyzer, with samples meeting a minimum RIN of 7.0 advanced to library preparation. Libraries were generated with the Illumina TruSeq Stranded RNA Kit with Ribo-Zero Gold and sequenced on NovaSeq (Illumina) instruments at 100PE.

Sequencing reads were trimmed of primers and poly-A tails using Cutadapt [56]. The trimmed sequences were then aligned to a combined human GRCh38 and mouse GRCm38 reference genome with STAR [57]. The alignment and resulting counts table obtained from STAR was split into mouse and human tables based on Ensemble IDs and then used for downstream analyses.

### Consensus Molecular Subtype Classification

Human-mapped RNA-seq raw counts from 24 3D-eCRC-PDX tissues and CRC-PDX tumors were combined with 157 colon adenocarcinoma PDX datasets from the NCI PDMR [58] to produce a unified counts matrix for consensus molecular subtype analysis. CMS classification was conducted using *CMScaller* [59] under default settings for RNA-seq data. *CMScaller* applies a nearest template prediction approach [60], computing cosine distances between each sample and defined CMS templates to assign the closest subtype [61]. Samples with prediction FDR values > 0.05 remained unclassified.

### Differential gene expression analysis with TCGA data

HT-Seq counts for colorectal adenocarcinoma tumors and matched tumor-adjacent tissues were downloaded from the TCGA-COAD [62] dataset through *TCGAbiolinks* [63]. These data were combined with HT-Seq counts generated from CRC-PDX tumors and engineered 3D-eCRC-PDX tissues by aligning all datasets using Ensembl gene IDs. Differential expression analysis was conducted using *DESeq2* [64], applying internal normalization and variance modeling. Significantly regulated genes were identified based on an FDR < 0.05. Genes meeting the thresholds of log_2_ fold change > 1.5 and baseMean > 10 were retained for subsequent pathway and gene set analyses.

### Gene Ontology Enrichment Analysis

Enrichment analyses for biological process and molecular function categories were conducted using the *clusterProfiler* R package [65]. Differentially expressed genes derived from two pairwise comparisons—CRC-PDX tumors relative to TCGA CRC tumor-adjacent tissue and 3D-eCRC-PDX tissues relative to TCGA CRC tumor-adjacent tissue—were used to identify overrepresented gene ontology terms and functional pathways.

### GSEA Analysis

Gene Set Enrichment Analysis (GSEA) was performed using *clusterProfiler* [65] with the Hallmark [66] and C4 Computational gene sets from the MSigDB database. The C4 Computational gene sets were defined by mining large collections of cancer-oriented expression data [67–69]. Three subcollections make up the C4 Computational gene sets: Curated Cancer Cell Atlas (3CA), Cancer gene neighborhoods (CGN), and Cancer modules (CM). Results were visualized using bubble plots, which display the normalized enrichment score (NES) and corresponding false discovery rate (FDR) q-value, providing an estimate of the likelihood that the enrichment represents a false-positive finding.

### Statistical Analysis

Hierarchical clustering was conducted using the *heatmap.plus* package implemented in RStudio (Version 1.2.1335). All subsequent statistical analyses were performed in GraphPad Prism 8.0 (GraphPad Software, San Diego, California USA, www.graphpad.com). Outliers were removed using the ROUT method (Q = 1%). Normality of the data distribution and homogeneity of variance were assessed prior to statistical testing. Depending on the experimental design, either one-way or two-way ANOVA followed by Tukey *post hoc* testing was used to evaluate group differences. Slopes representing rates of change over time were derived from linear regression and compared using an F-test. Statistical significance was defined as p ≤ 0.05 unless otherwise stated.

## Results

### Viable Cell Numbers Isolated from CRC-PDX Tumors were Patient Line-dependent

We previously established a tissue engineering platform to maintain CRC-PDX cells in culture [37]. Here we use our tissue engineering platform to examine colon cancer patient-to-patient heterogeneity using three CRC PDX lines from stage II, III-B, IV patient tumors (Fig. 1). We first determined the number of viable cells from the three different PDX lines and found that the number of viable cells isolated from CRC-PDX tumors was dependent on the patient PDX line (Suppl. Fig. 1). CRC-PDX tumors from the stage III-B patient yielded the highest cell number (35.9 ± 5.3 million per gr tumor), followed by stage IV (27.1 ± 5.3 million per gr tumor), and then stage II (15.2 ± 1.7 million per gr tumor) (*p* < 0.0208). The cells were encapsulated in PEG-Fb hydrogels and maintained their viability over time (Suppl. Fig. 2).

**Figure 1.**
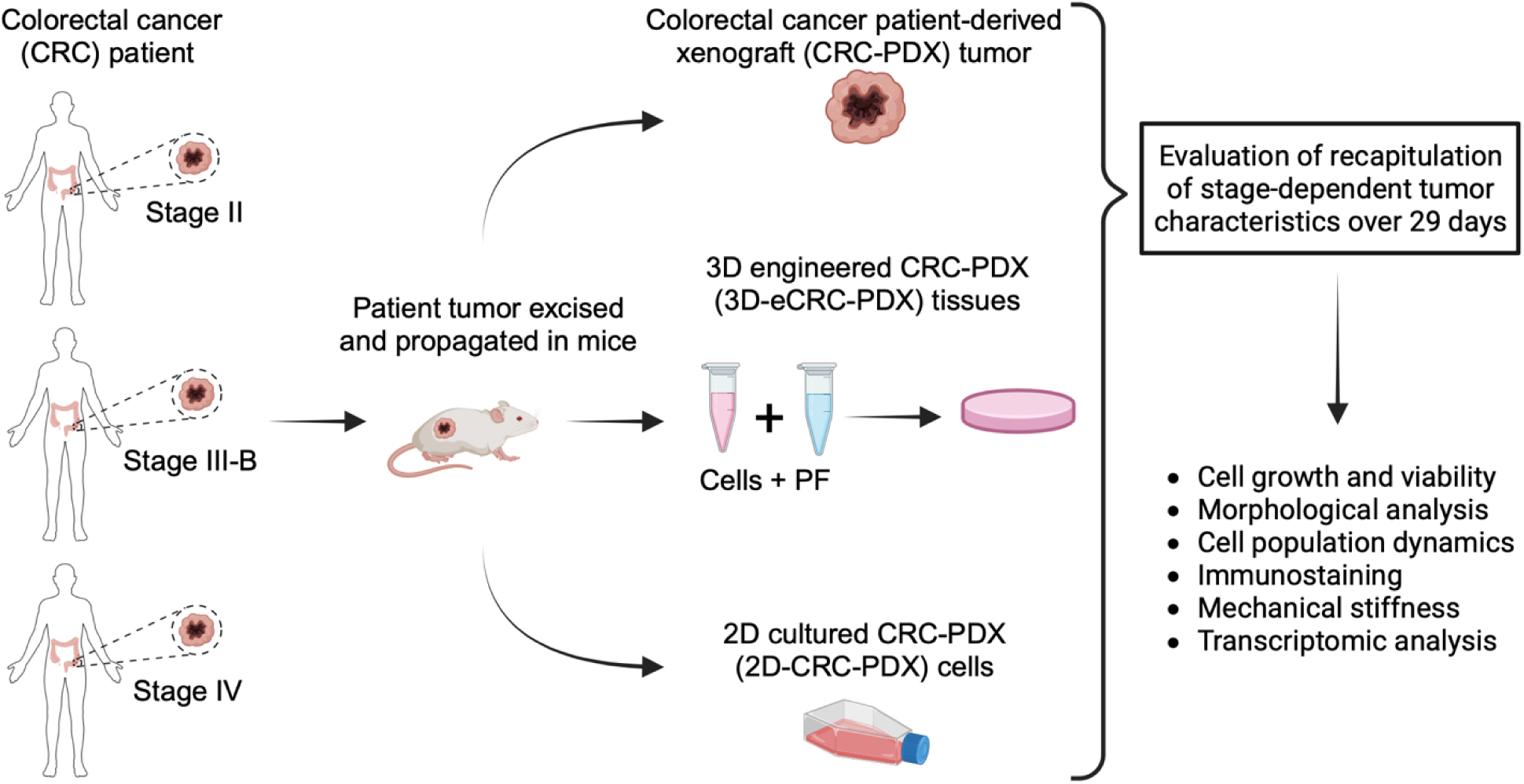
Tissue-engineered colorectal cancer models were established to recapitulate *in vivo* CRC-PDX tumor properties. Recapitulation of key CRC-PDX tumor properties was assessed in engineered tissues formed using CRC-PDX tumor cells and 2D cultured CRC-PDX cells during long-term *in vitro* culture. Patient-derived xenograft (PDX) tumors from three colorectal cancer (CRC) patients (referred to here as Stage II, Stage III-B, and Stage IV) were propagated in mice. These CRC-PDX tumors were characterized, and the cells dissociated from the tumors were then cultured in both 3D tissues (referred to as 3D-eCRC-PDX) and 2D cultures (referred to as 2D-CRC-PDX). Tests were conducted to assess the capability of the 3D-eCRC-PDX tissues and 2D-CRC-PDX cell cultures to recapitulate the differing characteristics of the originating CRC-PDX tumors.

### Temporal Growth and Morphological Changes of Tumor Cell Colonies Were Both PDX Line- and Culture Platform-dependent

Long-term in vitro culture models, including 3D engineered CRC-PDX tissues (3D-eCRC-PDX tissues) and 2D cultured cells (2D-CRC-PDX culture), were formed from CRC-PDX tumor cells, and the ability to maintain patient line-specific CRC-PDX tumor properties was evaluated (Fig. 1). To assess the temporal growth and morphological changes of cancer cell colonies in the 3D-eCRC-PDX tissues and 2D-CRC-PDX cell cultures, phase-contrast Z-stacks and images were taken over 29 days of cell culture (Fig. 2 A, B). For each CRC-PDX tumor line, three separate batches, each from a separate tumor, were prepared and analyzed. The analyzed data from all three batches were pooled for each CRC-PDX tumor line and are presented in Fig. 2 (see Suppl. Fig. 3, Suppl. Fig. 4, and Suppl. Fig. 5 for data from each batch). The data showed that the changes in area and diameter of cell colonies were both PDX line- and culture platform-dependent (Fig. 2 C-J and Suppl. Fig. 6). Within the 3D-eCRC-PDX tissues, the area and diameter of both stage II and stage III-B CRC-PDX cell colonies increased significantly from Day 1 to Day 29 (*p* < 0.0001); however, the rates of increase in area and diameter of stage II cell colonies were approximately 60-fold and 16-fold higher than those of stage III-B cell colonies, respectively (*p* < 0.0001) (Fig. 2 C, E and Suppl. Fig. 6 A, B). Interestingly, these significantly higher increases in colony and diameter of cell colonies for stage II 3D-eCRC-PDX tissues were consistent with the *in vivo* growth rates of CRC-PDX tumors (Fig. 2 C, E, Suppl. Fig. 6 A, B and Suppl. Fig. 7). On the other hand, the area and diameter of stage IV cell colonies increased significantly from Day 1 to Day 8 and then decreased thereafter (*p* < 0.0001) (Fig. 2 C, E and Suppl. Fig. 6 A, B).

**Figure 2.**
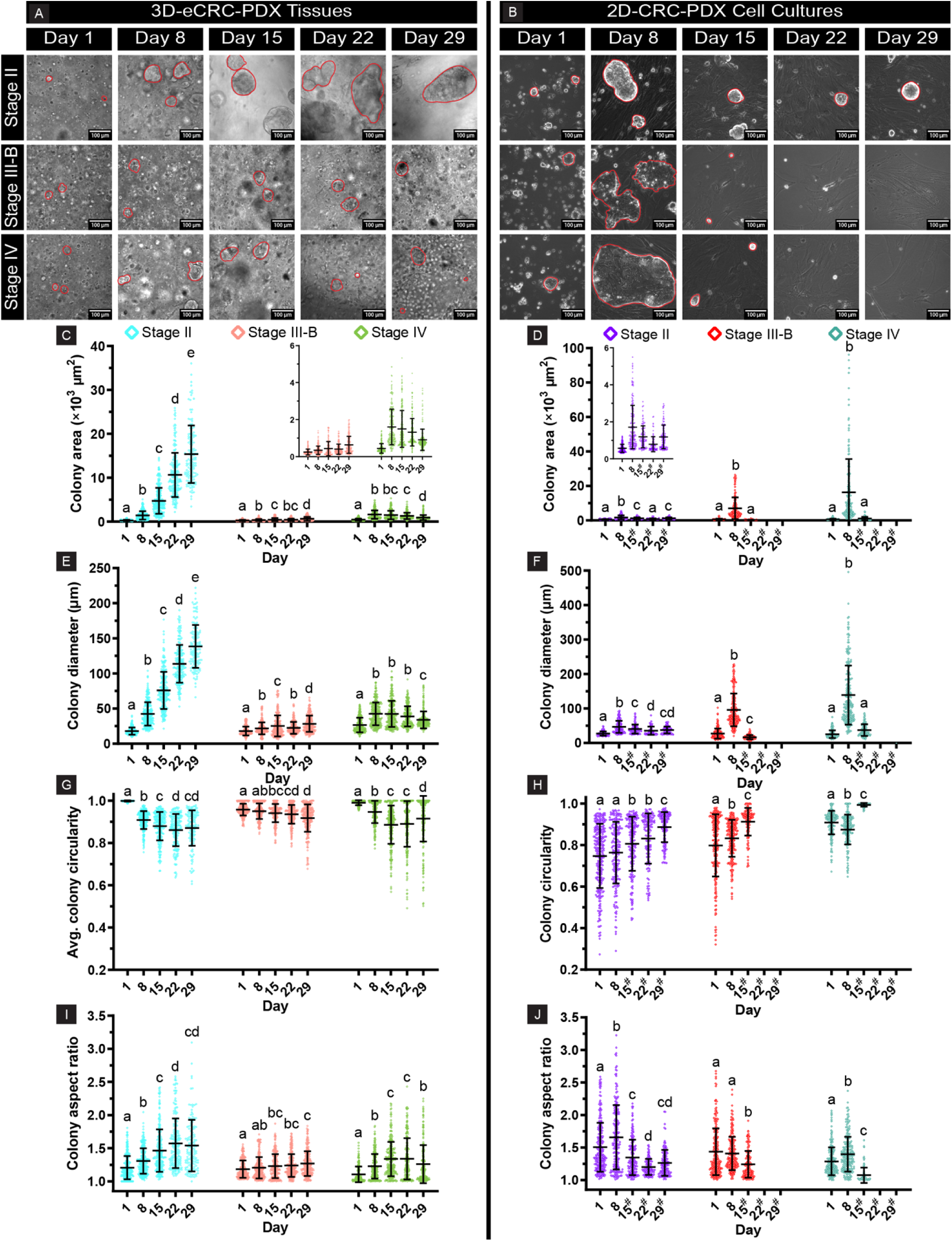
Growth and morphological changes of tumor cell colonies were both line- and culture platform-dependent. Representative phase-contrast images of cell colonies (outlined) in the (A) 3D-eCRC-PDX tissues and (B) 2D-CRC-PDX cell cultures over 29 days for all CRC lines. 3D-eCRC-PDX tissue (C) colony area and (E) colony diameter. Relative differences between lines in 3D-eCRC-PDX tissue recapitulate tumor growth rates (Supplemental Fig. 7), whereas 2D-CRC-PDX cell culture (D) colony area and (F) colony diameter did not. (Following Day 15 cell passaging, colonies did not re-form in stage III-B and stage IV 2D-CRC-PDX cell cultures). In 3D-eCRC-PDX tissues, generally, (G) colony circularity decreased, and (I) colony aspect ratio increased over time. In 2D-CRC-PDX cell cultures, generally, colony (H) circularity increased, and (J) aspect ratio decreased over time. All data are shown as mean ± SD, with each point representing one colony. A minimum of 80 colonies per CRC-PDX tumor line per time point per culture platform were evaluated for each of the three independently generated batches. Statistical significance was assessed across time points for the same cell line, and groups with different letters are significantly different. The # symbol indicates that 2D-CRC-PDX cells were passaged at the preceding time point.

For 2D-CRC-PDX cells, passaging was performed on Day 8 of in vitro culture; without passaging on Day 8, cell numbers plateaued and did not increase thereafter (Suppl. Fig. 8). In 2D-CRC-PDX cell cultures, the area and diameter of CRC-PDX cell colonies increased significantly from Day 1 to Day 8 for all CRC-PDX tumor lines (*p* < 0.0001); however, stage IV CRC-PDX cells demonstrated the largest colony area and diameter, followed by stage III-B, and then stage II (*p* < 0.0001) (Fig. 2 D, F). Following cell passaging, although the colonies re-formed in stage II 2D-CRC-PDX cell cultures over 29 days, the resulting colonies remained smaller than those quantified on Day 8. In both stage III-B and stage IV 2D-CRC-PDX cell cultures, only a few colonies re-formed after the first cell passaging (from Day 8 to Day 15), and no colonies re-formed thereafter.

Cell colony circularity and aspect ratio were also assessed over 29 days of culture. Within the 3D-eCRC-PDX tissues, the circularity of both stage II and stage III-B CRC-PDX cell colonies decreased and the aspect ratio of those increased significantly from Day 1 to Day 29 (*p* < 0.0001); however, the rates of change for stage II and stage III-B CRC-PDX cell colonies were significantly different (*p* < 0.0001) (Fig. 2 G, I and Suppl. Fig. 6 C, D). The rate of decrease in circularity and increase in aspect ratio of stage II cell colonies was approximately 2.4-fold and 3.9-fold higher than those of stage III-B cell colonies, respectively. On the other hand, the circularity of stage IV cell colonies decreased and aspect ratio increased significantly from Day 1 to Day 15 (*p* < 0.0001); after Day 15, the circularity increased and aspect ratio decreased (*p* < 0.0045) (Fig. 2 G, I and Suppl. Fig. 6 C, D). For 2D CRC-PDX cultures, stage II cell colonies showed significant increasing and decreasing trends for circularity and aspect ratio, respectively, from Day 1 to Day 29 (*p* < 0.0001) (Fig. 2 H, J). Similar to the stage II CRC-PDX tumor line, the stage III-B exhibited increasing and decreasing trends for circularity and aspect ratio, respectively, from Day 1 to Day 15 (*p* < 0.0001) (no colonies re-formed after Day 15), whereas the circularity of stage IV cell colonies decreased and the aspect ratio increased from Day 1 to Day 8 (*p* < 0.005) with opposite trends from Day 8 to Day 15 (*p* < 0.0001).

Overall, both 3D-eCRC-PDX tissues and 2D-CRC-PDX cell cultures for all PDX lines showed similar growth patterns and morphological changes between three separate batches (Suppl. Fig. 3 and Suppl. Fig. 4); thus, these models demonstrated a high degree of reproducibility. However, in sharp contrast to 2D-CRC-PDX cell cultures, 3D-eCRC-PDX tissues were able to reproduce the PDX line-specific *in vivo* CRC-PDX tumor growth rates.

### 3D-eCRC-PDX Tissues Better Maintained Cell Subpopulations as Compared to 2D-CRC-PDX Cell Cultures

As PDX tumors contain both human-derived and mouse-derived cells, maintaining these cell subpopulations in *in vitro* cultures is vital for recapitulating the *in vivo* tumors. To assess the ability of both 3D-eCRC-PDX tissues and 2D-eCRC-PDX cultures to maintain different cell subpopulations of the originating CRC-PDX tumors in *in vitro* cultures over 29 days, flow cytometry was used. To prepare single cells for flow cytometry cell quantification, we first dissociated and counted the viable cells in both 3D-eCRC-PDX tissues and 2D-CRC-PDX cell cultures at each time point using trypan blue and a hemocytometer. The cell counts showed that the rates of change in viable cell number were different between 3D-eCRC-PDX tissues and 2D-CRC-PDX cell cultures and were dependent on each CRC-PDX tumor line (Fig. 3 A, B). In 3D-eCRC-PDX tissues, the viable cell number increased significantly for all stages of CRC-PDX cells (*p* < 0.05); however, the rate of increase for stage II 3D-eCRC-PDX tissues was significantly higher than those for stage III-B and stage IV 3D-eCRC-PDX tissues (approximately 3.4-fold and 17.5-fold, respectively, *p* < 0.0001). Interestingly, these rates of increase in 3D-eCRC-PDX tissues were consistent with *in vivo* growth rates of the originating CRC-PDX tumors (Fig. 3 A and Suppl. Fig. 7). In sharp contrast, the rate of increase in viable cell number from Day 1 to Day 15 for stage III-B 2D-CRC-PDX cell cultures was significantly higher than those for stage II and stage IV (2.1-fold and 2.0-fold, respectively, *p* < 0.0165), with no significant difference between stage II and stage IV (Fig. 3 B). The stage II 2D-CRC-PDX cell number continued to increase over 29 days, whereas the cell numbers in stage III-B and stage IV 2D-CRC-PDX cell cultures decreased from Day 15 to Day 29 due to cell death.

**Figure 3.**
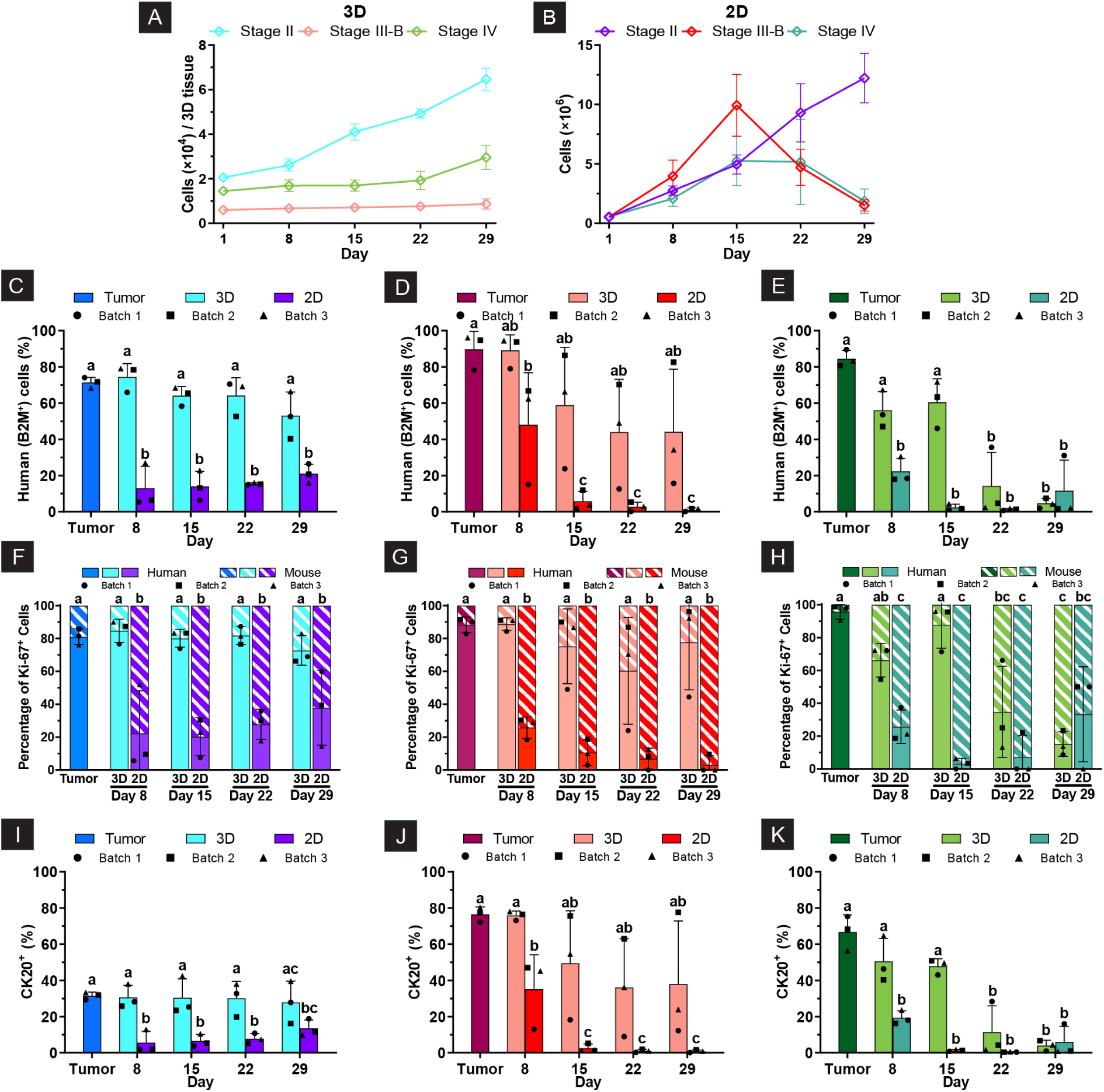
3D-eCRC-PDX tissues better maintained CRC-PDX tumor cell subpopulations as compared to 2D-eCRC-PDX cell cultures. (A) In 3D-eCRC-PDX tissues, the increase in viable cell number was significantly higher for stage II than for stage III-B and IV patient lines, matching in vivo CRC-PDX tumor growth rates (Suppl. Fig. 6). (B) In contrast, in 2D-CRC-PDX cell cultures, the viable cell number increased the most for stage III-B from Day 1 to Day 15, with no significant difference between stage II and IV. Stage II cell numbers continued increasing for 29 days, while stage III-B and IV cell numbers decreased after Day 15. (C-E) Human (B2M+) cell percentages were 71 ± 3% (stage II), 90 ± 10% (stage III-B), and 85 ± 5% (stage IV). In 3D-eCRC-PDX tissues, stage II and III-B maintained these subpopulations through day 29, while stage IV retained them for 15 days. In sharp contrast, in 2D-CRC-PDX cell cultures, human cell subpopulations decreased significantly after 8 days. (F-H) Stage II and III-B 3D-eCRC-PDX tissues maintained proliferative human cells (Ki67+, B2M+) for 29 days, while stage IV maintained them for 15 days. In sharp contrast, a significant decrease in this cell subpopulation was found after 8 days in 2D-CRC-PDX cell cultures, with percentages remaining low thereafter. (I-K) CK20 marker trends were similar to those of human cell populations. Data are shown as mean ± SD; groups labeled with different letters are significantly different (p ≤ 0.05, n = 3 separately generated batches of cultured engineered tissues or cells, with each batch originating from a separate CRC-PDX tumor).

To quantify the percentage of human-derived cells (human cells) and mouse-derived cells (mouse cells), B2M and H2Db markers were used, respectively. The human cell subpopulation percentages in CRC-PDX tumors were found to be PDX line-dependent. Approximately 71 ± 3%, 90 ± 10%, and 85 ± 5% of all cells were human cells for stage II, stage III-B, and stage IV CRC-PDX tumors, respectively (Fig. 3 C-E). Interestingly, the stage II and III-B 3D-eCRC-PDX tissues maintained this cell subpopulations over 29 days of culture; no significant difference was found between the percentages of human cell subpopulation in the 3D-eCRC-PDX tissues at each time point and the originating CRC-PDX tumor cells (*p* > 0.0649) (Fig. 3 C, D). The stage IV 3D-eCRC-PDX tissues were able to maintain the human cell subpopulations over 15 days of culture (*p* > 0.0551); however, the human cell subpopulation decreased by over 60% on Days 22 and 29 as compared to the originating CRC-PDX tumors in this line (Fig. 3 E). In sharp contrast, in 2D-CRC-PDX cell cultures, the percentages of human cell subpopulations decreased significantly after 8 days of culture from 71 ± 3% to 13 ± 12% for stage II, from 90 ± 10% to 48 ± 29% for III-B, and 85 ± 5% to 22 ± 7% for stage IV (*p* < 0.0001) (Fig. 3 C-E). In stage II and stage IV 2D-CRC-PDX cell cultures, these percentages remained constant after Day 8, whereas in stage III-B 2D-CRC-PDX cell cultures, the percentage of human cells decreased significantly from Day 8 to Day 15 (from 48 ± 29% 6 ± 5%, *p* < 0.05) and remained constant thereafter. It should be noted that the 3D-eCRC-PDX tissues remained in uninterrupted culture for 29 days, whereas the 2D-CRC-PDX cells necessitated passaging every 7 days beginning on Day 8 to avoid confluence (Suppl. Fig. 8). Data was collected from 2D-CRC-PDX cells prior to each passage.

To explore the mechanism underlying the superior preservation of human cell subpopulations in 3D-eCRC-PDX tissues relative to 2D-CRC-PDX cultures, we quantified proliferative populations by co-staining Ki-67 with B2M or H2Db to distinguish human from mouse proliferating cells. In stage II CRC-PDX tumors, 23 ± 4% of the total cells were Ki-67+, and 81 ± 5% of these proliferating cells were human (Fig. 3 F and Suppl. Fig. 9 A). Notably, the stage II 3D-eCRC-PDX tissues maintained this proliferative human cell subpopulations over 29 days of culture, whereas in 2D-CRC-PDX cells, a significant decrease in this cell subpopulation was found after 8 days of culture (from 81 ± 5% to 22 ± 26%, *p* = 0.0002) with the percentages remaining low thereafter (Fig. 3 F). Similar to stage II 3D-eCRC-PDX tissues, no significant difference was found between the percentages of proliferative human cell subpopulation in the stage III-B 3D-eCRC-PDX tissues and originating CRC-PDX tumors (88 ± 5%, *p* > 0.2571), whereas stage III-B 2D-CRC-PDX cell cultures showed significantly lower percentages on all time points as compared to that for the originating CRC-PDX tumors (*p* < 0.0013) (Fig. 3 G and Suppl. Fig. 9 B). For stage IV CRC-PDX tumors, although only 6 ± 2% of cells were proliferative, 95 ± 4% of these proliferative cells were human cells (Fig. 3 H and Suppl. Fig. 9 C). The stage IV 3D-eCRC-PDX tissues maintained this proliferative human cell subpopulation over 15 days; however, a significant decrease in the percentages of this subpopulation was observed on Day 22 (from 95 ± 4% to 35 ± 28%, *p* < 0.0008) and Day 29 (from 95 ± 4% to 15 ± 7%, *p* < 0.0001) (Fig. 3 H). Similar to stage II and stage III-B 2D-CRC-PDX cell cultures, the percentage of proliferative human cell subpopulation in stage IV 2D-CRC-PDX cell cultures decreased significantly after 8 days of culture (from 95 ± 4% to 26 ± 10%, *p* < 0.0001) and remained low thereafter (Fig. 3 H).

To investigate the temporal changes in CRC cell subpopulations, cytokeratin 20 (CK20) was used in tandem with B2M or H2Db. We previously (CITE) showed that not all CRC cells express CK20, and CK20^−^ cells may include other CRC cell subpopulations. Stage II, stage III-B, and stage IV CRC-PDX tumors had 32 ± 2%, 77 ± 4%, and 67 ± 10% CK20^+^ cells, respectively (Fig. 3 I-K). Both stage II and stage III-B 3D-eCRC-PDX tissues maintained this cell subpopulation over 29 days of culture (Fig. 3 I, J), whereas stage IV 3D-eCRC-PDX tissues maintained it over 15 days of culture with a significant decrease on Day 22 (from 67 ± 10% to 11 ± 15%, *p* < 0.0001) and Day 29 (from 67 ± 10% to 4 ± 3%, *p* < 0.0001) (Fig. 3 K). In sharp contrast, this cell subpopulation decreased significantly from Day 1 to Day 8 in all stages of 2D-eCRC-PDX cultures (from 32 ± 2% to 6 ± 6% for stage II, 77 ± 4% to 35 ± 19% for stage III-B, and 67 ± 10% to 19 ± 4% for stage IV, *p* < 0.05) and remained low thereafter (Fig. 3 I-K). These trends were also observed for the dual-labeled B2M^+^ (or H2Db^−^) and CK20^−^ cell subpopulations (Suppl. Fig. 9 D-F). In addition, as expected for all stages of CRC, the percentages of CK20^+^ and human CK20^+^ cell subpopulations were equivalent (Fig. 3 I-K and Suppl. Fig. 9 G-I), meaning that the CK20^+^ cells were a subpopulation of the human cells while all mouse cells were B2M^−^ (or H2Db^+^) as well as CK20^−^.

Overall, 3D-eCRC-PDX tissues consistently maintained the proportions of key cell subpopulations—remaining unchanged for 29 days in the stage II and III-B models and for 15 days in the stage IV model—closely reflecting the originating CRC-PDX tumors. In contrast, 2D-CRC-PDX cultures showed significant shifts after only 8 days. For key cell subpopulations within the 3D-eCRC-PDX tissues, batch-to-batch variabilities were found to be higher for the stage III-B PDX line as compared to stage II and IV lines; thus, the stage II and IV 3D-eCRC-PDX tissues were more reproducible.

### 3D-eCRC-PDX Tissue Immunostaining Identifies PDX Line-dependent Regional Distribution of Human and Stomal Cells

Whole tissue staining for F-actin and CK20 showed the distribution of the colonies of CK20+ cells and stromal cells over time (Days 15-29, Suppl. Fig. 10). To more clearly visualize the stage II, III-B, and IV 3D-eCRC-PDX human and stromal cell populations in the center and at the edge of the 3D-eCRC-PDX tissues, day 15 tissues were cryosectioned and immunostained (Suppl. Fig. 11A). Human cells were found to be distributed as colonies in both the center and at the edge of the stage II 3D-eCRC-PDX tissues (Suppl. Fig. 11B) as visualized through human mitochondria staining (MAB1273, Clone 113-1 antibody) but not at the edge in the stage III-B and IV 3D-eCRC-PDX tissues (Suppl. Fig. 11B). As visualized by WGA staining to label cellular membrane glycoproteins, non-human (mouse-derived) stromal cells were found to be localized within colonies of human cells and distributed throughout the tissue in both the center and at the edge in the stage II 3D-eCRC-PDX tissues (Suppl. Fig. 11B). In stage IV 3D-eCRC-PDX tissues, these stromal cells were also present within human cell colonies at the center. However, in stage III-B 3D-eCRC-PDX tissues, they were not localized within the human cell colonies, instead exhibiting a more widespread distribution throughout the tissue center (Suppl. Fig. 11B). At the edge of the stage III-B and IV 3D-eCRC-PDX tissues, staining indicated that the stromal cells were also more widely distributed (Suppl. Fig. 11B). These findings suggest that the stage III-B and stage IV 3D-eCRC-PDX tissues have a greater stromal component than the stage II 3D-eCRC-PDX tissues. However, the distribution of the stromal component was found to be PDX line dependent. These findings are consistent with the cell subpopulation analysis observed using flow cytometry.

### 3D-eCRC-PDX Tissues Reproduced PDX Line Specific Differences in Mechanical Stiffness of CRC-PDX Tumors

To quantify tissue stiffness, parallel plate compression testing was performed on at least two independent CRC-PDX tumors and two paired 3D-eCRC-PDX tissue batches for each CRC stage. Stiffness measurements revealed significant regional differences within CRC-PDX tumors for all stages; the periphery of each CRC-PDX tumor showed the highest stiffness, followed by the midpoint, and then the tumor core (*p* < 0.05) (Fig. 4 A-C). CRC stage PDX line-dependent differences in stiffness were also observed. The average tumor stiffness of all batches (pooled data from all tumors of a given line) was found to be PDX line dependent. Upon closer examination, line-to-line differences were driven by geometric region-specific disparities. Although the average stiffness values for the core were equivalent for all CRC stages (0.49 ± 0.08, 0.46 ± 0.06, and 0.41 ± 0.07 kPa for stage II, III-B, and IV, respectively, *p* > 0.8849), the periphery of stage II CRC-PDX tumors had the highest stiffness (2.95 ± 0.55 kPa), followed by those of stage III-B (1.94 ± 0.54 kPa), and then stage IV (1.24 ± 0.27 kPa) (*p* < 0.0001) (Suppl. Fig. 12 A). Furthermore, the consistency of stiffness between batches (tumors) of a given CRC stage was also found to be PDX line dependent. Whereas the stiffness values of a given region were equivalent for batch 1 and batch 2 of stage II CRC-PDX tumors (*p* = 0.9928) (Fig. 4A), significant variations between batches were observed for both midpoint and periphery of both stage III-B and stage IV CRC-PDX tumors (*p* < 0.0338); no significant difference between batches was found for the core (*p* > 0.8253) (Fig. 4 B, C) for any line.

**Figure 4.**
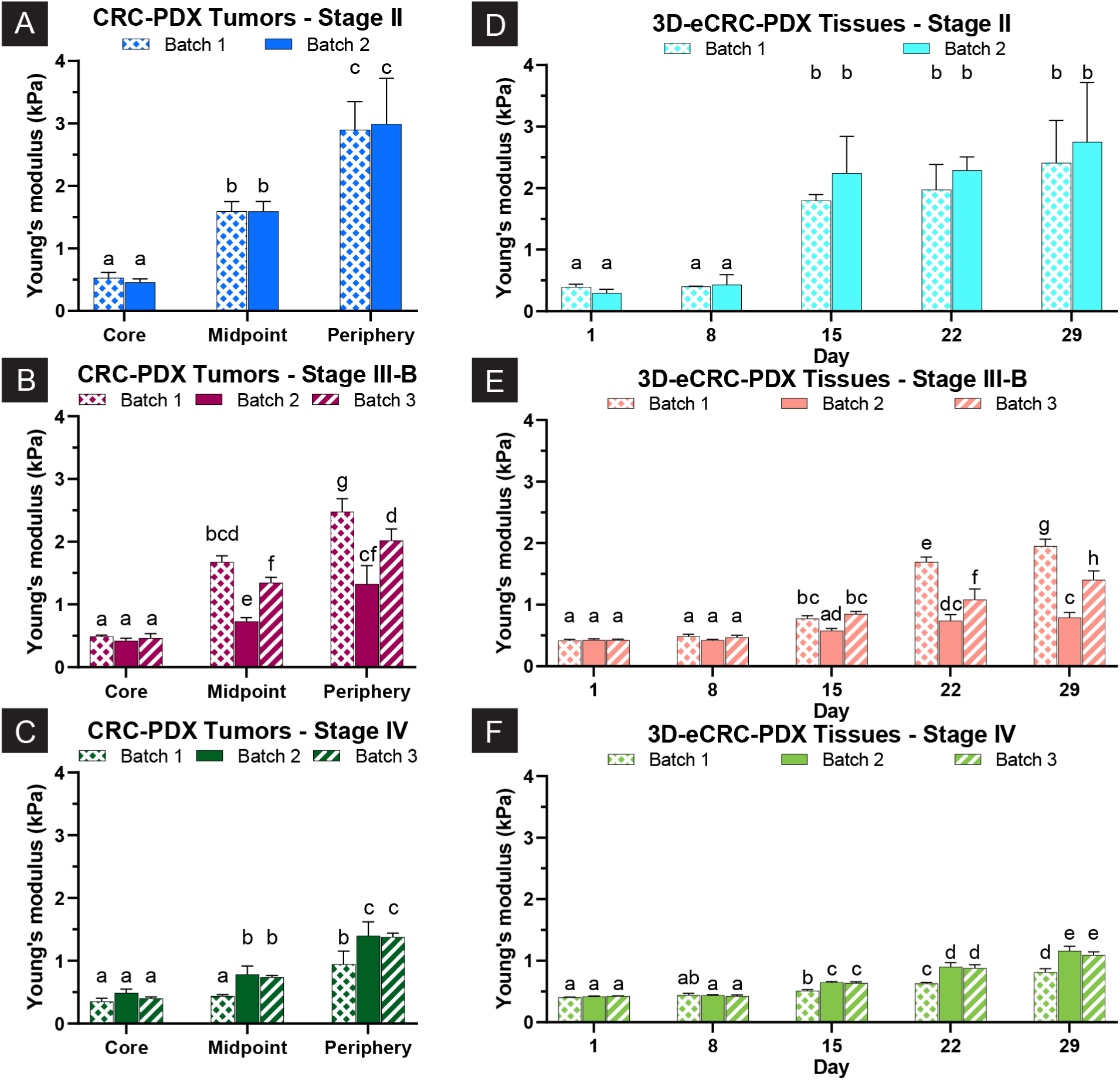
3D-eCRC-PDX tissues mimicked the mechanical stiffness of CRC-PDX tumors. (A-C) For all CRC stages, CRC-PDX tumor stiffness varied regionally with an increasing trend from the tumor core to the periphery. Whereas the stiffness values of a given region were equivalent for batch 1 and batch 2 of (A) stage II CRC-PDX tumors, significant variations among batches were observed for both the midpoint and periphery of both (B) stage III-B and (C) stage IV CRC-PDX tumors; no significant difference among batches was found for the core. (D-F) The stiffness of 3D-eCRC-PDX tissues increased significantly over time for all CRC stages. Time-dependent stiffness of the 3D-eCRC-PDX tissues mimicked both region-dependent and batch-dependent stiffness of the originating CRC-PDX tumors for a given CRC stage (D vs. A, E vs. B, and F vs. C). Data are shown as mean ± SD; groups labeled with different letters are significantly different. (*p* ≤ 0.05, n = 3 tissues per region or time point per batch).

The stiffness of 3D-eCRC-PDX tissues increased significantly over time (*p* < 0.0001) for all CRC stages (Fig. 4 D-F). Remarkably, for each CRC stage PDX line, the time-dependent stiffness of the 3D-eCRC-PDX tissues mimicked both region-dependent and batch-dependent stiffness of the originating CRC-PDX tumors (Fig. 4 D vs. A, E vs. B, and F vs. C). In addition, the average stiffness of all batches of 3D-eCRC-PDX tissues from a given line recapitulated the line-dependent average stiffness of the CRC-PDX tumors (Suppl. Fig. 12 A vs. B). Whereas the average stiffness values for 3D-eCRC-PDX tissues on Day 1 were equivalent for all CRC stages (0.35 ± 0.07, 0.42 ± 0.02 and 0.42 ± 0.01 kPa for stage II, III-B, and IV, respectively, *p* > 0.8681), on Day 29, stage II CRC-PDX tumors showed the highest stiffness (2.58 ± 0.78 kPa), followed by stage III-B (1.38 ± 0.51 kPa), and then stage IV (1.02 ± 0.17 kPa) (*p* < 0.0258) (Suppl. Fig. 12 B).

Since changes in mechanical stiffness can arise from ECM remodeling, we investigated the changes in size and shape of the 3D-eCRC-PDX tissues over 29 days for at least 2 separate batches of 3D-eCRC-PDX tissues per line. The analyzed data from all batches were pooled for each CRC-PDX tumor line and are presented in Fig. 5 (see Suppl. Fig. 13 for data from each batch). It was found that for all stages of CRC, the volume, diameter, and aspect ratio (side view) of the 3D-eCRC-PDX tissues decreased, and circularity (side view) increased significantly from Day 1 to Day 29 (*p* < 0.0001); however, the rates of change were CRC-PDX tumor line-dependent (Fig. 5 B-E). For instance, the rates of decrease in volume, diameter, and aspect ratio of stage II 3D-eCRC-PDX tissues were significantly higher (approximately 50%, 35%, and 30% for volume, diameter, and aspect ratio, respectively, *p* ≤ 0.05) than those of stage III-B and stage IV 3D-eCRC-PDX tissues (Fig. 5 B, C, E); the rate of increase in circularity of stage II 3D-eCRC-PDX tissues was significantly higher (60%, *p* < 0.0001) than those of stage III-B and stage IV 3D-eCRC-PDX tissues (Fig. 5 D). The rates of change in volume, diameter, circularity, and aspect ratio of stage III-B and stage IV 3D-eCRC-PDX tissues were equivalent (*p* > 0.1971) (Fig. 5 B-E). The changes in circularity and aspect ratio of the 3D-eCRC-PDX tissues were consistent with the changes of tissues from the initial disk-like shape to a spherical shape (Fig 5 A).

**Figure 5.**
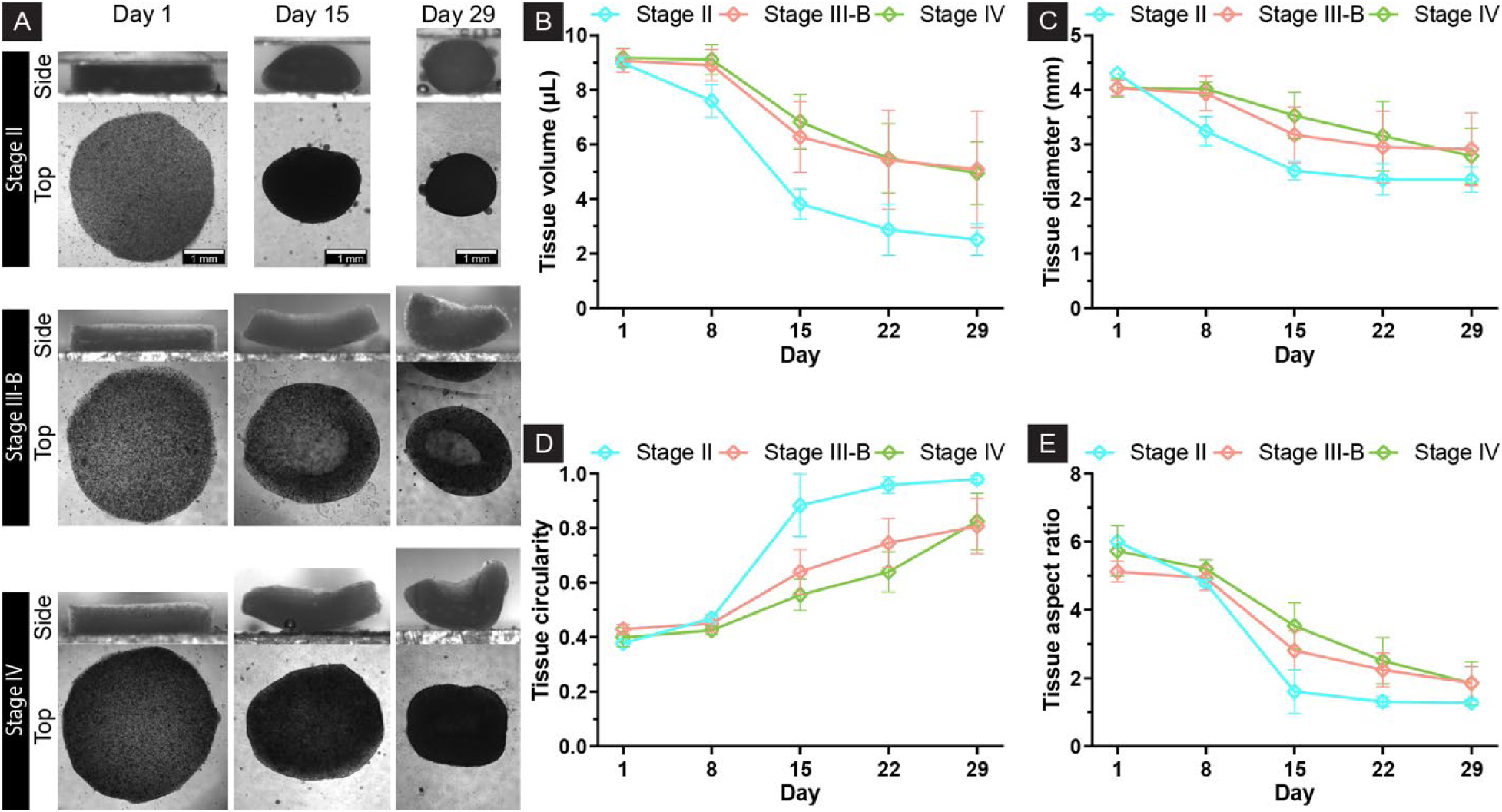
3D-eCRC-PDX Tissues Exhibited Cell-induced Contraction. (A) Representative phase-contrast images of the 3D-eCRC-PDX tissues over 29 days for all CRC stages. The rates of decrease in (B) volume, (C) diameter, and (E) aspect ratio (side view) and increase in (D) circularity (side view) of stage II 3D-eCRC-PDX tissues were significantly higher than those of stage III-B and stage IV 3D-eCRC-PDX tissues. The rates of change in volume, diameter, circularity, and aspect ratio of stage III-B and stage IV 3D-eCRC-PDX tissues were equivalent. The observed decrease in volume, diameter, and aspect ratio, together with the increase in circularity, corresponded to the transformation of 3D-eCRC-PDX tissues from disk-like to rounded morphologies. Data are expressed as mean ± SD. (*p* ≤ 0.05, n = a minimum of 6 tissues from at least 2 separate batches of cell culture).

### Transcriptomic Analysis Reveals Similarities and Differences between 3D-eCRC-PDX Tissues and CRC-PDX Tumors

To elucidate the molecular mechanisms underlying the maintenance of patient-to-patient PDX line heterogeneity in our tissue-engineered platform, we performed an RNA-seq transcriptomic analysis and mapped the data to the human and mouse genomes, assessing similarities and differences between CRC-PDX tumors and 3D-eCRC-PDX tissues after 15 days of in vitro culture. The greatest correspondence in human and mouse sequence read alignment was observed between the stage II 3D-eCRC-PDX tissues and CRC-PDX tumors (Suppl. Table 1).

We first sought to assess the molecular subtype [61] of the 3D-eCRC-PDX tissues and CRC-PDX tumors. Using RNA seq data from a cohort of 157 colon adenocarcinoma PDX tumors from the NCI Patient-Derived Models Repository (PDMR) in which 113 tumors were classified as CMS1-4, we classified the consensus molecular subtype of the 3D-eCRC-PDX tissues, CRC-PDX tumors, and PDMR PDX lines. Consistent with prior classifications of CRC tumors [61, 70], the majority of PDMR PDX tumors were classified as consensus molecular subtype (CMS) 2 and 4 (Suppl. Fig. 14A). We observed that two of the three stage II 3D-eCRC-PDX tissues and all three stage II CRC-PDX tumors classified as CMS1 (Suppl. Fig. 14B). In contrast, three stage III-B 3D-eCRC-PDX tissues classified as CMS4, while three of the five stage III-B CRC-PDX tumors classified as CMS2. Similar findings were observed in the stage IV 3D-eCRC-PDX tissues and CRC-PDX tumors: tissues classified as CMS4 and tumors classified as CMS2 (Suppl. Fig. 14B). These findings suggest CMS recapitulation on Day 15 is PDX line dependent.

We next sought to identify patient-specific similarities in human cancer-related genes in the CRC-PDX tumors and the 3D-eCRC-PDX tissues by performing an analysis using RNA seq data from normal colon tissue and the CRC-PDX tumors and then separately with normal colon tissue and the 3D-eCRC-PDX tissues. We observed that stage II 3D-eCRC-PDX tissues and CRC-PDX tumors had the highest percentage of overlapping DEGs (68%) with normal colon tissues, while the stage IV 3D-eCRC-PDX tissues and CRC-PDX tumors had the fewest overlapping DEGs (51%) with normal colon tissues (Fig. 6A).

**Figure 6.**
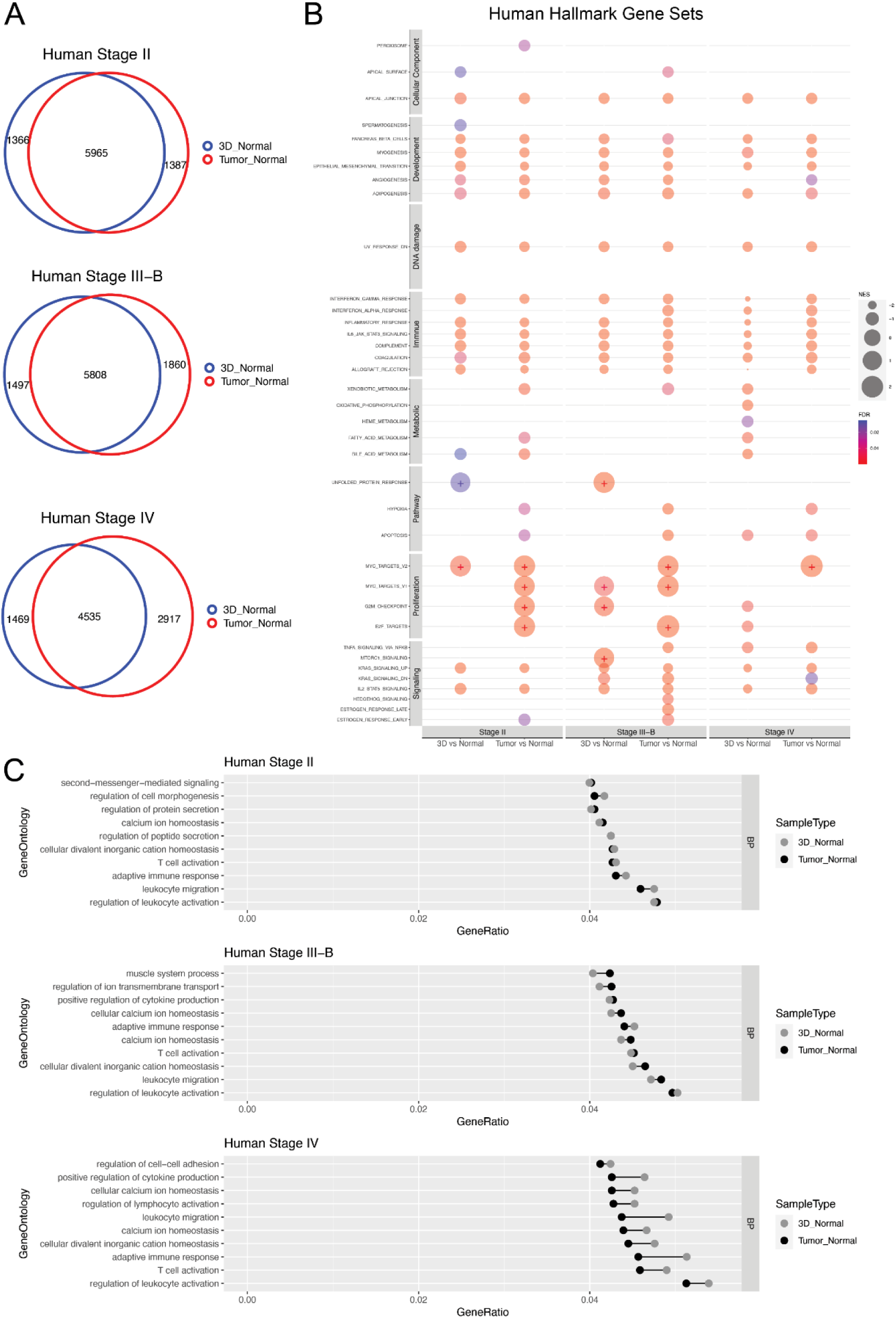
Patient-dependent transcriptomic profiles were recapitulated in 3D-eCRC-PDX tissues. (A) DEGs in CRC-PDX tumors and 3D-eCRC-PDX tissues compared to normal colon tissues (n = 41) from the TCGA-COAD. The number of total and overlapping DEGs is shown for each stage. (B) GSEA analysis showing the top enriched Hallmark gene sets in CRC-PDX tumors and 3D-eCRC-PDX tissues compared to normal colon tissue (n = 41) for each stage. Positive enrichments are indicated using a “+” within the bubble. (C) Gene ontology analysis of biological process enriched in DEGs between CRC-PDX tumors and 3D-eCRC-PDX tissues and normal colon tissues (n = 41) for each stage.

To determine patient-specific similarities in cancer-related pathways, we examined enriched GSEA Hallmark gene sets in the CRC-PDX tumors and the 3D-eCRC-PDX tissues compared to normal colon tissue (Fig. 6B). When examining enrichment for all the Hallmark gene sets, directionality of gene set enrichment was consistent in all comparisons. Similar enrichment for Hallmark gene sets related to cellular component, development, DNA damage, and immune response between normal colon tissue and both 3D-eCRC-PDX tissues and CRC-PDX tumors was observed for all three stages. In particular, both 3D-eCRC-PDX tissues and CRC-PDX tumors from all stages showed highly consistent enrichment compared to normal colon tissue in KRAS signaling and IL2-STAT5 signaling gene sets. When examining individual gene sets within the metabolic, pathway, proliferation, and signaling groups of Hallmark gene sets, there was not consistent enrichment of all Hallmark gene sets for both 3D-eCRC-PDX tissues and CRC-PDX tumors as compared to normal colon tissue. Metabolic-related gene sets were enriched in stage IV 3D-eCRC-PDX tissues but not in their corresponding CRC-PDX tumors and not in stage II and III-B engineered tissues. In the pathway-related gene sets, unfolded-protein-response was highly enriched in 3D-eCRC-PDX tissues for stage II and III-B but not in other conditions; hypoxia-related gene sets were enriched in all CRC-PDX tumors but not in the 3D-eCRC-PDX tissues. To further examine patient-specific similarities in cancer-related pathways, we assessed enriched GSEA C4 Computational gene sets (large collections of cancer-oriented expression data [67–69]) from the human mapped RNA seq data in the CRC-PDX tumors and the 3D-eCRC-PDX tissues compared to normal colon tissue (Suppl. Fig. 15A). As was also seen with the Hallmark gene set enrichment, there was not consistent enrichment of all C4 Computational gene sets for both 3D-eCRC-PDX tissues and CRC-PDX tumors as compared to normal colon tissue.

To further assess patient-specific similarities in cancer-related pathways, we performed a gene ontology (GO) analysis to examine over-representation in the biological process (BP) category. The top 10 ranked GO BP terms according to gene ratio (percentage of total DEGs in the given GO term) of the 3D-eCRC-PDX tissues and CRC-PDX tumors vs normal colon tissues are shown (Fig. 6C). Although ranked differently, 3D-eCRC-PDX tissues and CRC-PDX tumors for each stage had the same GO BP terms in the top 10. Consistent with the DEG analysis, the greatest correspondence between the top 10 ranked GO BP terms in the 3D-eCRC-PDX tissues and CRC-PDX tumors was observed in the stage II PDX line, while the least correspondence was observed in the stage IV PDX line. Consistent with the DEG analysis and the top 10 ranked GO BP terms, the greatest correspondence between the GO molecular functions (MF) terms in the 3D-eCRC-PDX tissues and CRC-PDX tumors was observed in the stage II PDX line (Suppl. Fig. 16). Finally, to gain insight into patient-specific similarities in ECM related gene sets, we examined enriched GSEA curated matrisome gene sets [71, 72] in the CRC-PDX tumors and the 3D-eCRC-PDX tissues compared to normal colon tissue (Suppl. Fig. 17A). We observed negative enrichment in all 3D-eCRC-PDX tissues and CRC-PDX tumors compared to normal colon tissue for all matrisome gene sets (Suppl. Fig. 17A).

To determine patient-specific differences in cancer-related pathways, we next examined enriched GSEA Hallmark gene sets from the human mapped RNA seq data in the CRC-PDX tumors compared to the 3D-eCRC-PDX tissues (Fig. 7A). The stage III-B 3D-eCRC-PDX tissues show significant enrichment across all categories in the Hallmark gene sets compared to the stage II CRC-PDX tumors which had the lowest number of enriched Hallmark gene sets. Enrichment in the TNFA_Signaling_Via_NFKB gene set was the only gene set in which enrichment was observed in 3D-eCRC-PDX tissues from each PDX line. To further assess patient specific cancer-oriented gene expression differences in 3D-eCRC-PDX tissues compared to CRC-PDX tumors, we examined enrichment of GSEA C4 Computational gene sets. The stage IV 3D-eCRC-PDX tissues show significant enrichment across the most cancer-oriented computational genes compared to the stage II and III-B CRC-PDX tumors (Suppl. Fig. 15B).

**Figure 7.**
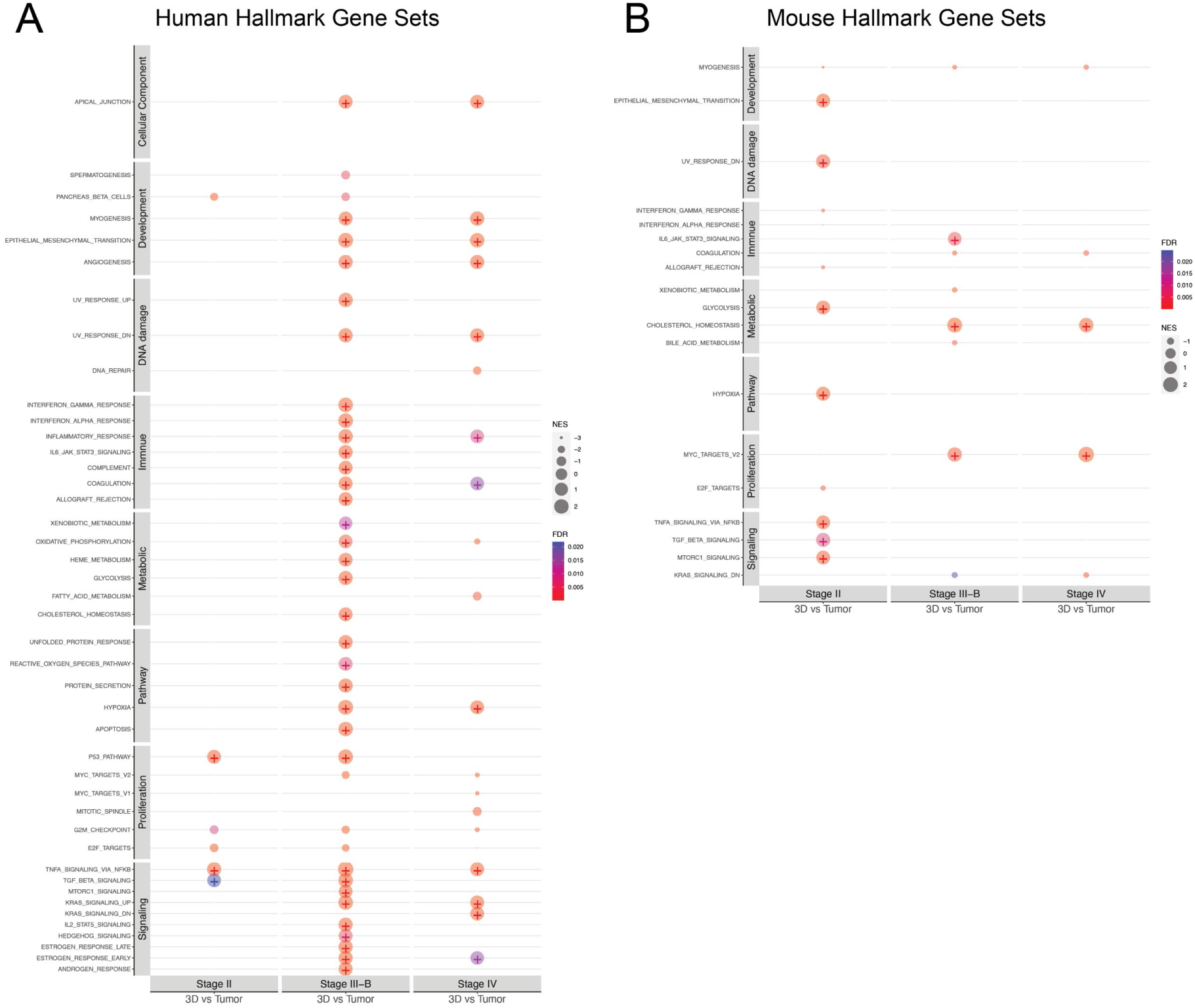
GSEA analysis between 3D-eCRC-PDX tissues and CRC-PDX tumors for human and mouse for each stage. GSEA analysis showing the top enriched Hallmark gene sets in 3D-eCRC-PDX tissues compared to CRC-PDX tumors for each stage for (A) human and (B) mouse. The bubble size indicates the normalized enrichment score (NES), and positive enrichment is labeled with a “+” within the bubble. The bubble color indicates the false discovery rate (FDR).

To gain insight into ECM-related gene sets, GSEA-curated matrisome gene sets [71, 72] were examined for enrichment in the 3D-eCRC-PDX tissues. We observed positive enrichment in the stage IV 3D-eCRC-PDX tissues for the NABA_Core_Matrisome gene set, while negative enrichment in the stage II 3D-eCRC-PDX tissues (Suppl. Fig. 17B). Similar enrichment findings were observed in additional matrisome gene sets: NABA_proteoglycans, NABA_ECM_regulators, NABA_ECM_glycoproteins, NABA_collagens (Suppl. Fig. 17B).

We next performed a transcriptomic analysis using the RNA seq data from the CRC-PDX tumors and 3D-eCRC-PDX tissues mapped to the mouse genome to determine whether the stromal component of 3D-eCRC-PDX tissues was enriched compared to the CRC-PDX tumors for GSEA Hallmark gene sets (Fig. 7B). Only the myogenesis gene set was enriched (negatively) in the 3D-eCRC-PDX tissues from all three PDX lines. Overlapping enrichment in gene sets was observed in stage III-B and stage IV 3D-eCRC-PDX tissues, although there were fewer enriched gene sets in the stage IV compared to III-B 3D-eCRC-PDX tissues (5 vs. 8, respectively). When the 3D-eCRC-PDX tissues from 3 different patients were compared to each other, we observed that murine-mapped gene sets in Proliferation (MITOTIC_SPINDLE, G2M_CHECKPOINT, and E2F_TARGETS) were positively enriched in the stage III-B and IV 3D-eCRC-PDX tissues compared to the stage II 3D-eCRC-PDX tissues but negatively enriched in the CRC-PDX tumors. The unique pattern of enriched gene sets from the murine-mapped RNA seq data suggests that PDX line-specific transcriptomic differences were maintained in the stromal component of the 3D-eCRC-PDX tissues.

## Discussion

To understand tumor biology and investigate disease mechanisms, *in vitro* CRC models that mimic patients’ tumors with high physiological relevance are needed. However, current *in vitro* models do not recapitulate patient-to-patient variabilities and tumor microenvironment heterogeneity with regard to tumor stiffness and stromal/fibroblast incorporation. PDX tumors, an *in vivo* model, have the ability to maintain parental tumor heterogeneity [11, 14–17]; however, the use of PDX tumors, as an animal model, is typically time-consuming, labor-intensive, and low throughput [18–20]. Here, we utilized CRC-PDX cells from three CRC patients (stage II, stage III-B, and stage IV CRC adenocarcinoma) in our previously developed models of 3D engineered CRC-PDX (3D-eCRC-PDX) tissues and 2D cultured cells (2D-CRC-PDX culture) and investigated the ability of our 3D and 2D models to recapitulate the patient-to-patient PDX tumor variabilities in comparison to the originating CRC-PDX tumors. We demonstrated that CRC-PDX cell colony sizes and cell numbers in 3D-eCRC-PDX tissues increased over time in a patient-specific manner similar to that of the originating CRC-PDX tumors, whereas cell numbers of 2D-CRC-PDX cell cultures did not. These 3D-eCRC-PDX tissues much better maintained cell subpopulations of the originating CRC-PDX tumors throughout long-term culture in comparison to 2D-CRC-PDX cell cultures. In addition, the mechanical stiffness of the 3D-eCRC-PDX tissues mimicked that of CRC-PDX tumors. Transcriptomic similarities and differences in 3D-eCRC-PDX tissues compared to the originating CRC-PDX tumors were observed in a PDX line-dependent manner. Taken together, our findings suggest that engineered tissues formed using CRC-PDX tumor cells can be employed and refined to build *in vitro* models that more closely recapitulate patient-specific CRC variability and tumor microenvironment heterogeneity.

To assess patient-specific similarities and differences in how tumor-derived cells behave in long-term 3D culture, we first examined growth kinetics across our three CRC-PDX–derived models. We found that the stage II 3D-eCRC-PDX tissues exhibited sustained and robust proliferation over 29 days, whereas the stage III-B and stage IV tissues showed reduced growth over the same period. The differential growth patterns observed among our three PDX models warrant careful interpretation. Importantly, maintaining PDX tumor cells in culture for 29 days, or even 15 days, represents an extraordinarily long culture period for these cells. PDX cells are notoriously difficult to maintain ex vivo for extended periods, and most studies are limited to much shorter timeframes [33, 48, 73, 74]. The supportive nature of our 3D platform is demonstrated by data showing maintenance of robust growth in the stage II model throughout 29 days and maintenance of cell viability within the stage III-B and IV models, even with limited proliferation. Importantly, these patterns mirror the in vivo behavior of the original PDX tumors (Supplementary Fig. 7): stage II grew robustly, while stage III-B and IV showed relatively limited proliferation. Although additional modulation of the platform-provided microenvironmental cues could potentially enhance proliferation of specific patient lines, the observed concordance indicates that our 3D-eCRC-PDX model preserves intrinsic, patient-specific tumor characteristics—particularly the altered growth dynamics and other microenvironmental dependencies common in advanced-stage disease—rather than necessarily reflecting a limitation of the platform. These results do indicate that there are PDX line-to-line differences in the optimal duration of in vitro culture prior to use of 3D-eCRC-PDX tissues for downstream analysis.

Cell-cell and cell-matrix interactions play a vital role in cell growth and proliferation and can affect changes in distinct cell subpopulations [21, 22, 75]. To mimic the cell-cell and cell-matrix interactions of a PDX tumor in an *in vitro* model, results suggest that all cell types from the originating PDX tumor should be incorporated in the *in vitro* tumor microenvironment. A few studies have reported the culture of PDX cells *in vitro* in a 3D matrix; however, these models did not include all cell types of the PDX cell population [33, 76]. Therefore, here we examined changes in distinct cell subpopulations using markers and investigated patient-to-patient tumor cell composition variabilities. Notably, we showed that the 3D-eCRC-PDX tissues maintained the key cell subpopulations of the originating stage II and stage III-B CRC-PDX tumors in long-term culture; these key cell subpopulations were also maintained in the stage IV 3D-eCRC-PDX tissues over at least 15 days of culture. This maintenance of distinct cell subpopulations occurred simultaneously with an increase in the total cell number in all stages of 3D-eCRC-PDX tissues. In sharp contrast, significant changes in all cell subpopulations occurred in 2D-CRC-PDX cell cultures. Kodack et al. [22] also reported a similar observation for 2D cultures of primary patient-derived tumors; they show that the stromal cells outgrew the cancer cells in 2D cultures, thereby impeding the cancer cell growth and resulting in the failure of cell line generation. In brief, our results demonstrated that the 3D-eCRC-PDX tissues for all stages mimicked the cell subpopulations of the originating CRC-PDX tumors, overcoming the limitations of other approaches.

Formation of our 3D-eCRC-PDX tissues required enzymatic and mechanical dissociation of PDX tumors into single cells, which disrupts their native 3D architecture. Therefore, here we assessed whether these dissociated cells could reorganize and re-establish key patient-specific tumor features through cell-autonomous behavior and cell–cell/matrix interactions when provided with a permissive PEG-Fb hydrogel supporting matrix. Although conceptually similar to organoid formation, our approach overcomes major organoid limitations: unlike organoid systems, the covalently crosslinked PEG-Fb hydrogel provides defined mechanics that support cell-mediated remodeling, extended culture (29 days), high cell density (20 million cells/cm^3^ initially), and cellular reorganization within the matrix to form a complete tissue rather than outward colony growth followed by subculture. Importantly, while organoid culture often loses stromal populations during passaging, our PDX-derived cultures retained both cancer and stromal cells throughout the culture period. Despite initial architectural disruption, PDX-derived cells rebuilt 3D structures exhibiting patient-specific growth kinetics, preserved heterogeneity, stiffness patterns reflective of the original tumors, and transcriptomic profiles that clustered with matched PDX tissue, with high batch-to-batch consistency. Thus, our model balances biological relevance with experimental control, offering a reproducible system for mechanistic studies.

The stiffness of tumor tissues has been demonstrated to have significant impacts on tumor cell phenotypes, such as proliferation, dormancy, migration, and metastasis. Therefore, stiffness is an important factor for cancer cell behavior [37, 52, 77–85]. Stiffness measurements were performed using parallel plate compression testing on intact 3D-eCRC-PDX constructs, without separating cells or cell colonies from the surrounding PEG-Fb hydrogel. This approach captures the composite contribution of both cellular aggregates and matrix, better reflecting the in vivo environment. CRC PDX tumor stiffness was similarly measured across multiple regions (core, midpoint, periphery) to account for spatial heterogeneity. While the Day 1 stiffness of 3D-eCRC-PDX tissues was similar across lines, it diverged over the 29-day culture period as cells remodeled the hydrogel, indicating that patient cell-specific matrix remodeling and tissue organization drive mechanical differences. Although PEG-Fb hydrogel mechanical properties are tunable, we used a standardized formulation across all three PDX lines to provide a consistent baseline environment. This approach allowed patient-specific differences in tumor mechanics to emerge through cell-mediated remodeling rather than variations in scaffold material. Importantly, the PEG-Fb scaffold undergoes degradation and remodeling over the 29-day culture period as cells modify their local environment, so the relative contribution of synthetic matrix versus cell-secreted ECM changes over time [51]. The observation of significant stiffness differences between the three PDX lines despite using identical starting PEG-Fb formulations indicates that cell-specific matrix remodeling and tissue organization drive the measured mechanical differences.

Although previous studies have not shown a comparison between *in vivo* and *in vitro* tissue stiffness, the stiffness of CRC tumors from patients is reported in the literature. For CRC samples directly taken from patients, higher tumor stiffness correlates with a higher stage, as well as metastatic phenotypes [86]. The bulk elastic modulus ranges of 1.91-3.96 kPa (median = 2.81 kPa), 1.08-12.2 kPa (median = 3.49 kPa), 2.60-43.2 (median = 8.89 kPa), and 5.58-68.0 kPa (median = 13.8 kPa) were observed for CRC of stages of I, II, III, and IV, respectively. However, both our CRC-PDX tumors and 3D-eCRC-PDX tissues demonstrated a reciprocal correlation between tissue stiffness and tumor stages. Possible reasons for this discrepancy between our result and those reported in literature could be the following: 1) the numbers of patients were small in both our studies and those in the literature; 2) the method of stiffness measurement used in the literature was not optimized; and 3) our PDX tumors were propagated subcutaneously whereas the literature reports the stiffness of the tumors taken directly from patients. Further research is warranted to compare CRC patient tumor, PDX tumor, and *in vitro* engineered tissue stiffness and the impact of initial *in vitro* engineered tissue stiffness on long-term *in vitro* tissue properties. Nonetheless, in our models, the average stiffness of all batches of 3D-eCRC-PDX tissues mimicked the stage-dependent average stiffness of the CRC-PDX tumors and was in the same order of magnitude as patient tumors [86].

We performed a transcriptomic analysis using RNA seq data to examine patient-to-patient similarities and differences between our CRC-PDX tumors and 3D-eCRC-PDX tissues. We observed that the stage II 3D-eCRC-PDX tissues and CRC-PDX tumors had the highest percentage of overlapping DEGs with normal colon tissues and the greatest correspondence between the top 10 ranked GO BP terms in the 3D-eCRC-PDX tissues and CRC-PDX tumors. In contrast, the least correspondence in DEGs with normal colon tissues and top 10 ranked GO BP terms in the 3D-eCRC-PDX tissues and CRC-PDX tumors was observed in the stage IV PDX line.

We also observed patient-to-patient differences between the CMS categorization of the CRC-PDX tumors and 3D-eCRC-PDX tissues for the stage III-B and IV PDX lines. The switch from a CMS2 canonical subtype [61] in the tumors to a CMS4 mesenchymal subtype [61] in the engineered tissues is consistent with our findings that the stage III-B and IV 3D-eCRC-PDX tissues had a greater percentage of mouse-mapped transcripts and more stromal cells detected at the edge of the 3D-eCRC-PDX tissues in immunostained tissue sections. These findings are also consistent with a positive enrichment in the stage IV 3D-eCRC-PDX tissues for the NABA_Core_Matrisome gene set and Metabolic-related gene sets. It is important to note that patient-derived organoids do not support the stromal CRC subtype (CMS4) [87]. Interestingly, we observed that compared the Stage III-B and IV PDX lines, the stage II PDX line had the greatest difference in mouse mapped transcripts between CRC-PDX tumors and 3D-eCRC-PDX tissues which both classified as the CMS1 subtype. This finding suggests that the CMS1 classification is less sensitive to changes in the stromal component in the CRC-PDX tumors and 3D-eCRC-PDX tissues.

Comparison of the 3D-eCRC-PDX tissues with existing 3D colorectal cancer (CRC) models underscores several strengths of our platform. Organoid-based systems are widely used and preserve epithelial characteristics while enabling genetic manipulation, but they frequently lack stromal components that are essential for capturing tumor microenvironment interactions [23–28]. Synthetic hydrogels such as PEG or PVA offer tunable mechanics and reproducible composition, yet they often lack endogenous biological cues needed for adhesion, migration, and matrix remodeling. Conversely, natural polymer scaffolds based on alginate, hyaluronic acid, or chitosan provide biocompatibility and cell-binding motifs but are difficult to precisely control in terms of degradation and mechanical properties. Hybrid materials aim to balance these limitations [88]. Microfluidic and bioprinted CRC models further enable spatial precision, perfusion, and construction of complex multicellular structures, though they typically rely on established cell lines and are designed for short-term studies, limiting their ability to model patient-specific heterogeneity or long-term tumor evolution [44, 89, 90]. Our PEG-Fb system addresses these gaps by integrating the tunability and consistency of a PEG-based synthetic matrix with the biological functionality of fibrinogen-derived motifs that support adhesion, spreading, and proteolytic remodeling [52]. Importantly, the use of PDX-derived cells allows for the long-term maintenance of both cancer and stromal populations—up to 29 days, which is notably extended for ex vivo PDX culture. During this period, cell-driven remodeling produces patient-specific stiffness profiles from identical initial formulations, effectively recreating each tumor’s microenvironment. The system also preserves transcriptomic alignment with the original PDX tumors and recapitulates in vivo growth patterns. By coupling reproducibility with biological fidelity, our study results support further development of this hybrid platform for mechanistic studies of tumor progression, stromal–cancer interactions, tumor evolution, and personalized therapeutic testing.

Although we demonstrated the ability of the 3D-eCRC-PDX tissues to mimic tumor-to-tumor variabilities, the current study had limitations. Our study serves as proof-of-concept that patient-to-patient heterogeneity can be preserved in the 3D-eCRC-PDX platform, and the batch-to-batch reproducibility demonstrated by the study establishes technical robustness. However, the limited number of PDX lines (n=3) constrains the generalizability of our findings. With only three lines representing stages II, III-B, and IV, we cannot draw definitive conclusions about stage-dependent differences or make systematic comparisons based on molecular subtypes. Broader biological conclusions regarding tumor subtype-specific behaviors require validation in a larger, molecularly characterized cohort. We are currently expanding our studies using more PDX lines with comprehensive molecular characterization, which will enable systematic comparison of tumor molecular subtypes. While our study demonstrated stage-dependent growth patterns in 3D-eCRC-PDX tissues, it provides an opportunity for future studies to build upon these findings through detailed quantitative analyses of fold-changes in cell number or colony volume across stages II, III-B, and IV. Additionally, further optimization of culture conditions—such as growth factor supplementation, nutrient adjustments, stromal-derived cytokines, or alternative matrix compositions—could enhance long-term proliferation and phenotypic maintenance of advanced-stage PDX cells. Although we were not able to include human stromal cells in our *in vitro* model as these cells are replaced by mouse stromal cells in PDX tumors, by using PDX tumor-derived stromal cells, we were able to assess RNA-seq data from human- and murine-mapped transcripts. Although employing CRC tumor cells dissociated directly from patient tumors to create 3D engineered tissues could lead to better tumor recapitulation, this approach would not have allowed us to evaluate batch-to-batch consistency when forming the 3D-eCRC-PDX tissues. Patient cells would have only allowed for the formation of one batch of engineered tissues, whereas by employing PDX-CRC tumors, we were able to evaluate consistency between batches when starting with a relatively consistent source of patient cells. Evaluation of *in vitro* culture platform consistency through batch-to-batch comparison, as performed here, is critical for the use of these models in downstream applications. Now that we have demonstrated the ability to maintain patient-to-patient PDX-CRC tumor heterogeneities, future work could directly utilize CRC patient tumor cells in 3D engineered tumor tissues to examine the ability to maintain patient tumor versus CRC-PDX tumor properties. Moreover, given previous work with other types of cancer cell lines in this model with and without stromal cell incorporation[52, 91, 92], our *in vitro* 3D model could be employed to examine the support of PDX or patient cells from other types of cancer.

## Conclusions

In this study, we employed three PDX tumor lines, originated from stages II, III-B, and IV tumors from CRC patients, encapsulated within PEG-Fb to examine to which extent the resulting engineered CRC-PDX tissues were able to recapitulate the variabilities between the originating CRC-PDX tumors. For all CRC stages, we demonstrated that the rates of change in CRC PDX cell colony sizes and cell numbers in 3D engineered CRC-PDX (3D-eCRC-PDX) tissues over time mimicked the growth rates of the originating CRC-PDX tumors. In sharp contrast to 2D cultured cells (2D-CRC-PDX culture), the 3D-eCRC-PDX tissues were able to maintain all cell subpopulations of the originating CRC-PDX tumors for all CRC stages. We monitored temporal variations in cell subpopulations in comparison to 2D-CRC-PDX cell cultures. Remarkably, the time-dependent stiffness of the 3D-eCRC-PDX tissues mimicked both region-dependent and batch-dependent stiffness of the originating CRC-PDX tumors for a given CRC stage. In addition, the average stiffness of all batches of 3D-eCRC-PDX tissues mimicked the stage-dependent average stiffness of the CRC-PDX tumors. Transcriptomic analysis revealed patient-specific similarities and differences in CRC-PDX tumors and 3D-eCRC-PDX tissues. The ability of these 3D-eCRC-PDX tissues to recapitulate the key attributes of the originating CRC-PDX tumors provides evidence that these engineered tissues are capable of mimicking patient-to-patient, underscoring the potential of our model to facilitate testing of personalized treatment strategies. In the future, this model can be refined and extended to include other types of cancer for investigation of disease mechanisms and use in drug testing applications.

## Data Availability

The RNA seq data sets (GSE151069 and GSE311262) are available from the NCBI Gene Expression Omnibus database (www.ncbi.nlm.nih.gov/geo). Flow cytometry data are available in the Supplementary Materials. All other data are available from the corresponding authors upon reasonable request.

## Acknowledgments

We sincerely thank Allison Church Bird at Auburn University’s Flow Cytometry Facility (Auburn, AL, USA) for her expert guidance in optimizing cell labeling procedures. We also appreciate the support of Dr. Mahmoud Mansour’s laboratory (Department of Anatomy, Physiology, and Pharmacology, Auburn University) for facilitating confocal microscopy experiments, and Dr. C. Ross Ethier’s laboratory (Wallace H. Coulter Department of Biomedical Engineering, Georgia Institute of Technology, Atlanta, GA, USA) for providing access to the MicroTester instrument. Some of the analyses in this work incorporate data generated by the TCGA Research Network (https://www.cancer.gov/tcga).

## Author Contributions

Conceptualization: M.J.H., M.W.G., E.A.L.; Methodology: I.H., B. Anbiah, N.L.H., Y.T., B. Ahmed, W.J.V., E.J.L., P.K., M.W.G., E.A.L.; Data collection and formal experimental analysis: I.H., B. Anbiah, N.L.H., B. Ahmed; Formal bioinformatics analysis: Y.T., W.J.V., E.J.L., M.W.G.; Investigation: I.H., B. Anbiah, N.L.H., Y.T., B. Ahmed, W.J.V., E.J.L., P.K., M.W.G., E.A.L.; Data Curation: I.H., N.L.H., Y.T., M.W.G., E.A.L.; Writing – original draft preparation: I.H., N.L.H., Y.T., W.J.V., E.J.L., M.W.G., E.A.L.; Writing – review and editing: I.H., B. Anbiah, N.L.H., Y.T., B. Ahmed, W.J.V., E.J.L., M.J.H., M.W.G., E.A.L.; Project administration: M.W.G., E.A.L.; Funding acquisition: E.J.L., M.J.H., M.W.G., E.A.L.

## Abbreviations

3D-eCRC-PDX: 3D Engineered Colorectal Cancer Patient-Derived Xenograft
AURIC: Auburn University Research Initiative in Cancer
B2M: Beta-2 Microglobulin
CK20: Cytokeratin 20
CMS: Consensus Molecular Subtype
CRC: Colorectal Cancer
DEGs: Differentially Expressed Genes
DMEM: Dulbecco’s Modified Eagle Medium
FBS: Fetal Bovine Serum
GO BP: Gene Ontology Biological Process
glutaGRO: GlutaGRO Supplement
GSEA: Gene Set Enrichment Analysis
H2Db: MHC Class I H-2 Db
IACUC: Institutional Animal Care and Use Committee
Ki-67: Ki-67 Proliferation Marker
MAB1273: Clone 113-1 Antibody
MSigDB: Molecular Signature Database
NABA: National Association for Biomedical Research
NCATS: National Center for Advancing Translational Sciences
NIH: National Institutes of Health
NIFA: National Institute of Food and Agriculture
NOD-SCID: Nonobese diabetic/severe combined immunodeficiency
PBS: Phosphate-Buffered Saline
PBS-T: Phosphate-Buffered Saline with Triton X-100
PDX: Patient-Derived Xenograft
PE: Phycoerythrin
PEG-Fb: Polyethylene Glycol-Fibrinogen
PEGDA: Poly(ethylene glycol) diacrylate
pen/strep: Penicillin-Streptomycin
RNA-seq: RNA Sequencing
TCGA: The Cancer Genome Atlas
WGA: Wheat Germ Agglutinin

