## Supplemental Figures for "Recapitulating Patient-to-Patient Colorectal Cancer Tumor Heterogeneity Using Patient-Derived Xenograft Cells in an Engineered Tissue Model"

cancer tissue engineering, PDX, colon cancer, organoid, stiffness, tumor microenvironment

**Supplementary Figures**

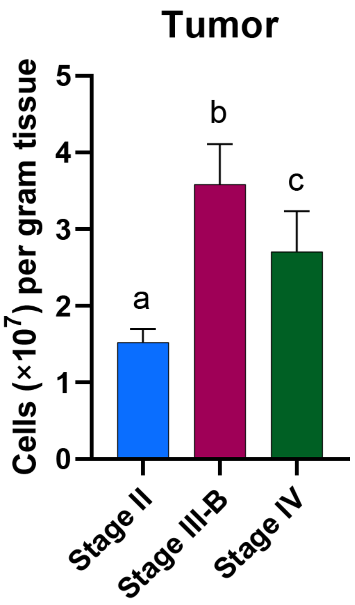


**Supplementary Figure 1. Viable cell numbers isolated from CRC-PDX tumors were line-dependent.** Viable cell number per gram of tumor was highest for stage III-B CRC-PDX tumors, followed by those for stage IV, and then stage II. Bar = mean ± SD. Means that do not share a letter are significantly different ($p < 0.03$; n = a minimum of 4 CRC-PDX tumors).

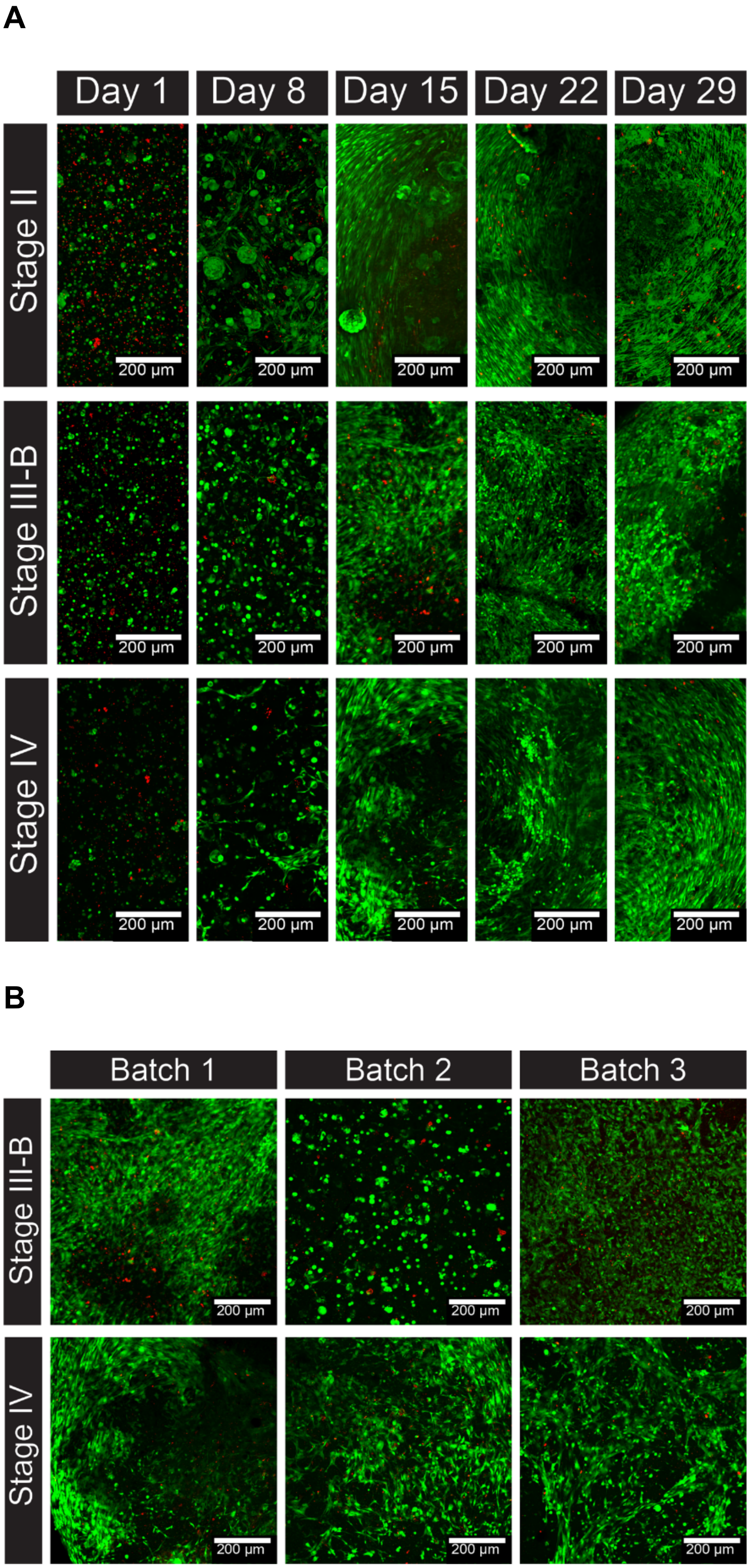

A
Day 1
Day 8
Day 15
Day 22
Day 29
Stage II
Stage III-B
Stage IV
200 μm
B
Batch 1
Batch 2
Batch 3
Stage III-B
Stage IV
200 μm

**Supplementary Figure 2. Representative confocal images for the viability assay**. (A) The images illustrate the presence of live (depicted in green) and dead (depicted in red) cells within 3D-eCRC-PDX tissues for all three CRC-PDX lines. Notably, there is observable cell elongation and an increase in cell colony size from day 1 to day 29 post-encapsulation. An increase in cell viability within the 3D-eCRC-PDX tissues was observed, potentially due to cellular proliferation. (B) The images underscore variability across three cell culture batches on Day 15, with a distinct observation in batch 2 for stage III-B indicating less pronounced cell elongation.

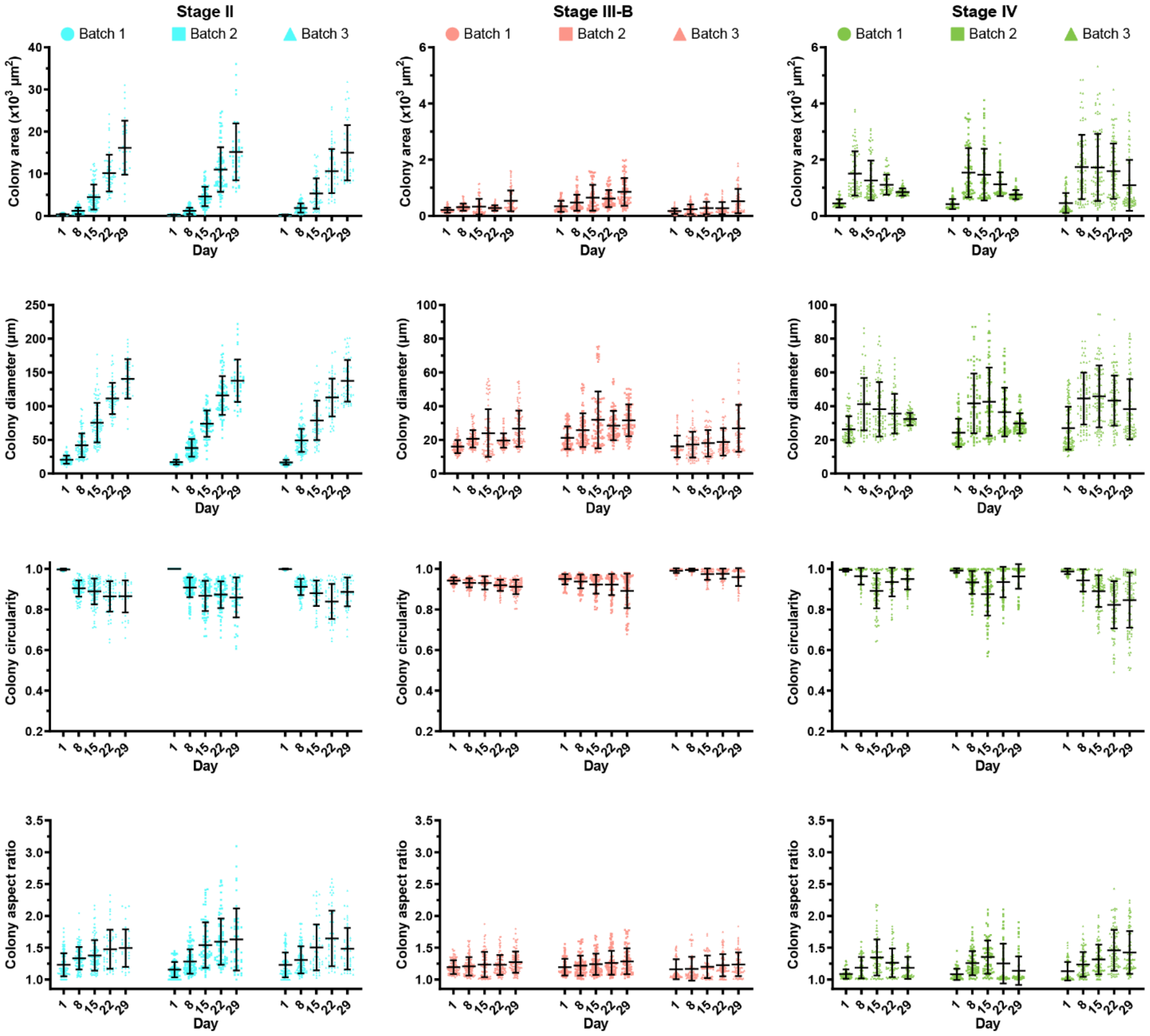


**Supplementary Figure 3. Quantification of growth and morphological changes of tumor cell colonies in 3D-eCRC-PDX tissues for 3 separate batches of cultures.** For each CRC stage, area, diameter, circularity, and aspect ratio of cell colonies were quantified over 29 days of culture. Data are mean ± SD, and each data point represents one colony. At least 80 CRC cell colonies per time point were examined for each of 3 separately prepared batches.

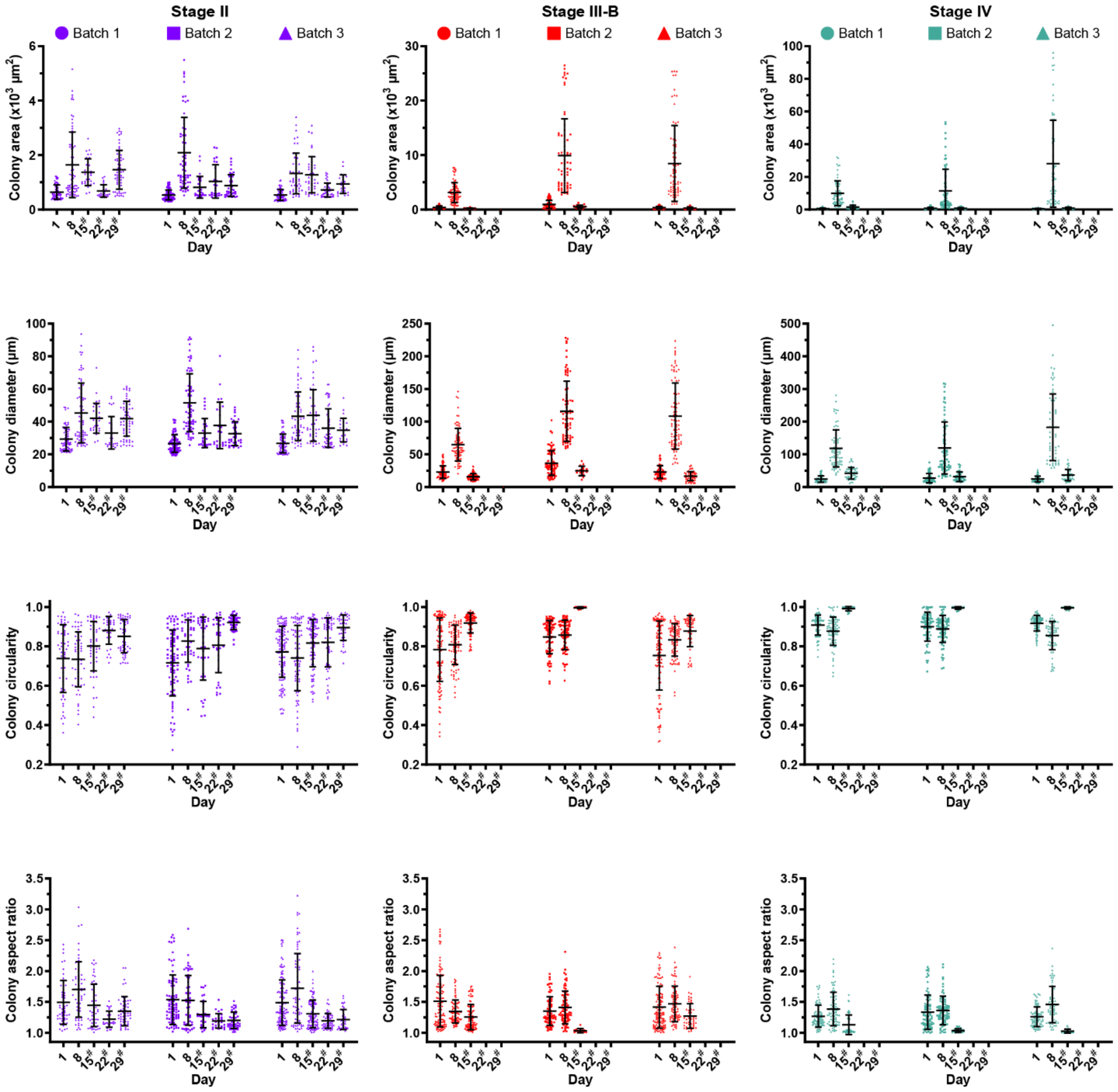


**Supplementary Figure 4. Quantification of growth and morphological changes of tumor cell colonies in 2D-CRC-PDX cultures for 3 separate batches of cultures.** For each CRC stage, area, diameter, circularity, and aspect ratio of cell colonies were quantified over 29 days of culture. Data are mean ± SD, and each data point represents one colony. At least 80 CRC cell colonies per time point were examined for each of 3 separately prepared batches. # indicates that the 2D-CRC-PDX cells were passaged at the previous time point.

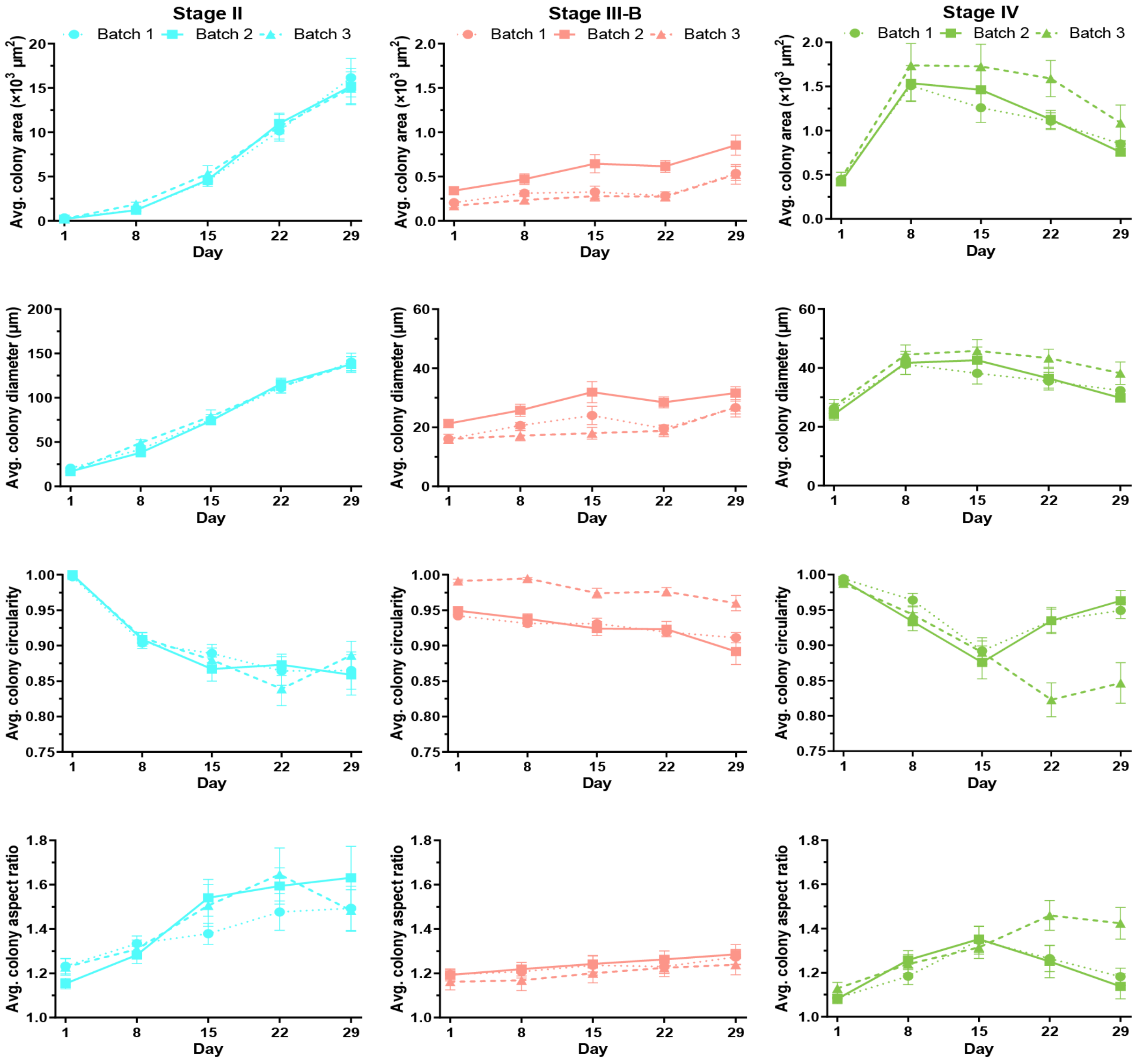


**Supplementary Figure 5. Quantification of growth and morphological changes of tumor cell colonies in 3D-eCRC-PDX tissues for 3 separate batches of cultures.** For each CRC stage, the averages of area, diameter, circularity, and aspect ratio of cell colonies were quantified over 29 days of culture. Data are mean ± SD and present the average values for at least 80 CRC cell colonies per time point.

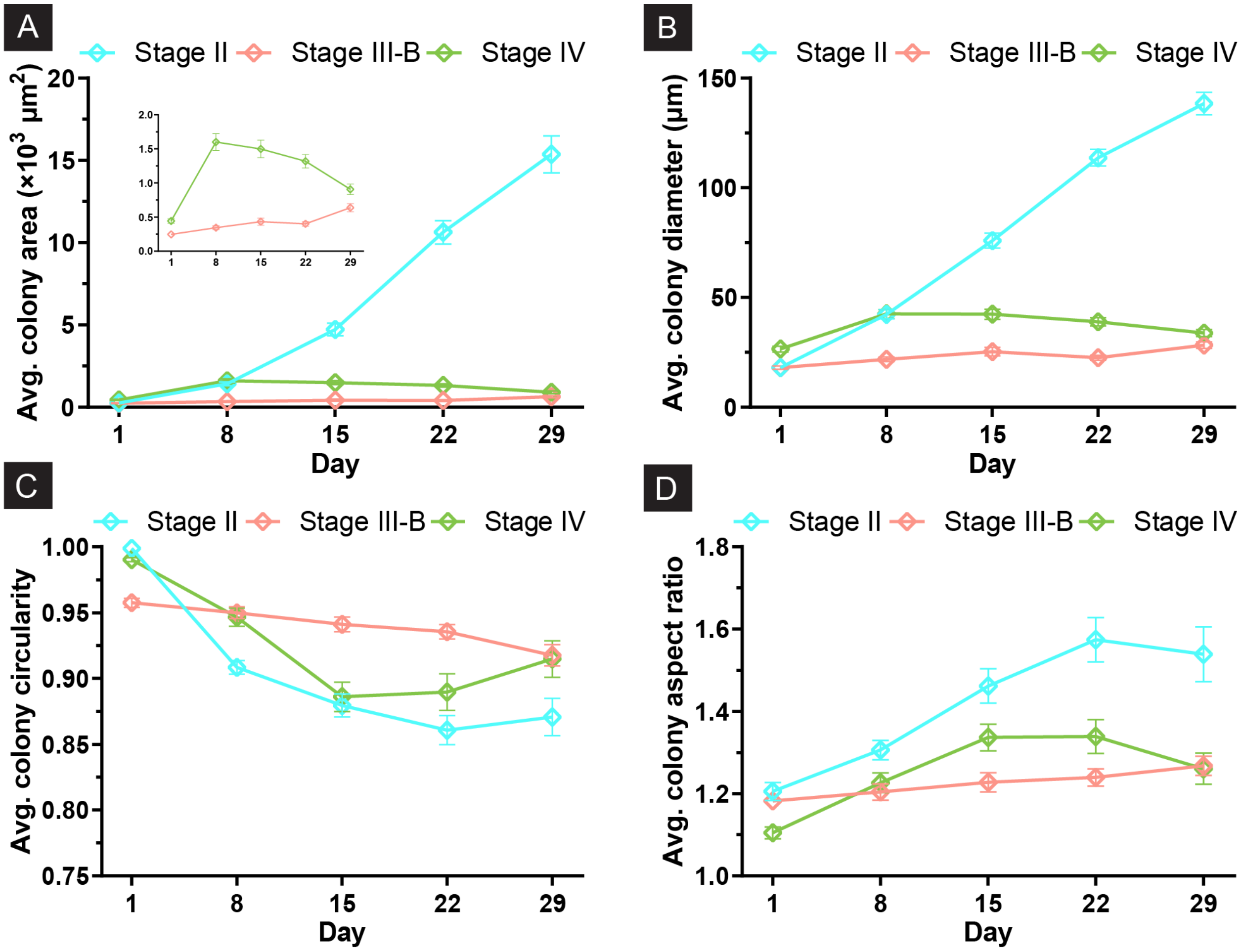


**Supplementary Figure 6. Quantification of growth and morphological changes of tumor cell colonies in 3D-eCRC-PDX tissues.** For each CRC stage, the averages of (A) area, (B) diameter, (C) circularity, and (D) aspect ratio of cell colonies were quantified over 29 days of culture. Data are mean ± SD and present the average values for at least 240 CRC cell colonies per time point, pooled from 3 separate batches of culture.

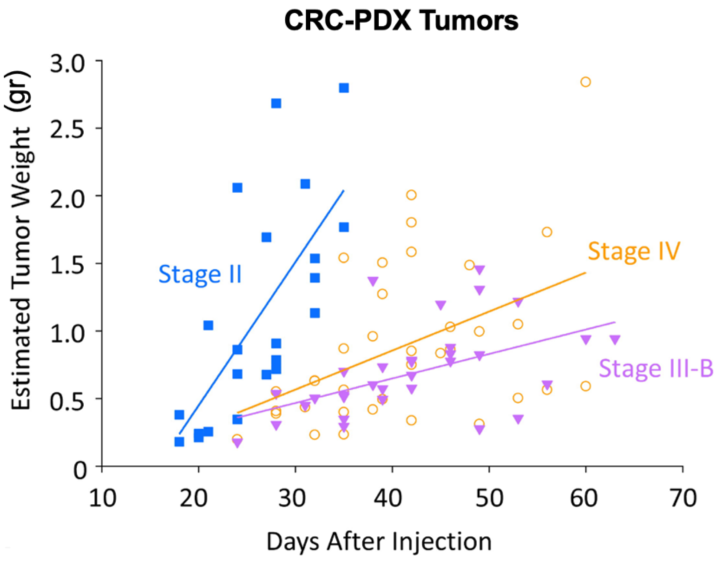


**Supplementary Figure 7. Quantification of CRC-PDX tumor growth.** For each CRC stage, the CRC-PDX tumors in mice were measured using a pair of calipers, and the estimated weight was calculated. Stage II CRC-PDX tumors showed a higher growth rate as compared to stage III-B and stage II CRC-PDX tumors (n= a minimum of 3-5 CRC-PDX tumors per line).

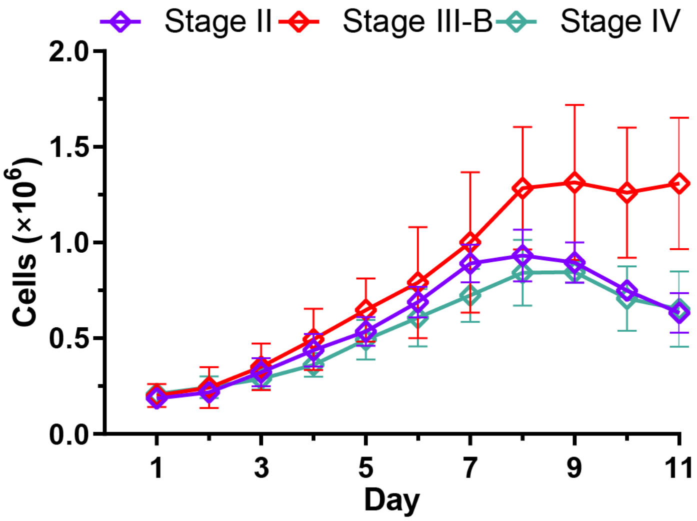


**Supplementary Figure 8. CRC-PDX cell growth rate in 2D-CRC-PDX cultures without cell passaging**. The total number of viable 2D-CRC-PDX cells, cultured in tissue culture flasks, increased from Day 1 to Day 8. An increase in cell number was not observed thereafter as cell confluency was reached on Day 8. Therefore, the 2D-CRC-PDX cells were passaged every 7 days for experiments comparing them to the 3D-eCRC-PDX tissues. Data are mean ± SD (n= 3 separate batches of cell cultures).

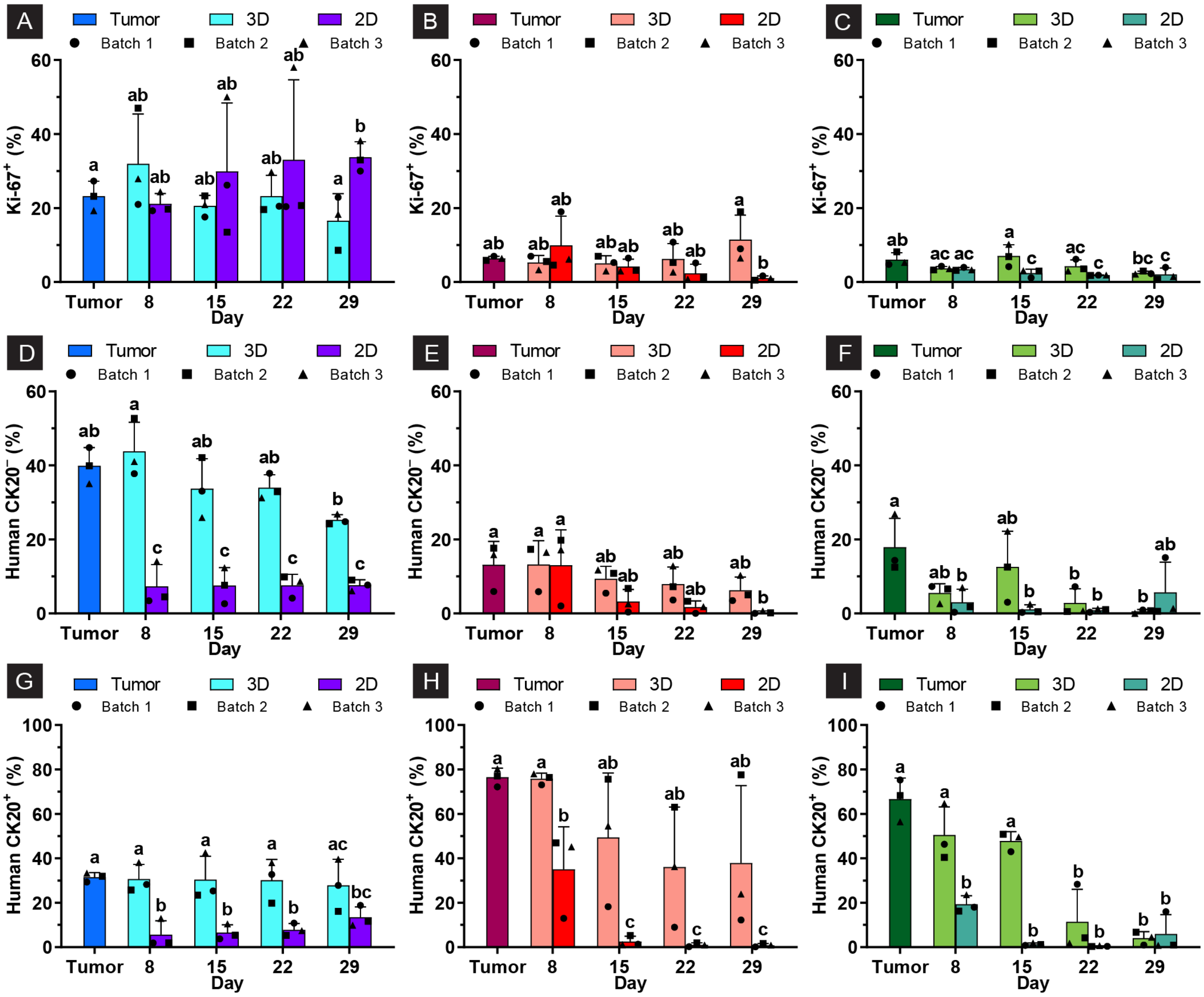

**Supplementary Figure 9. The 3D-eCRC-PDX tissues better maintained the originating CRC-PDX tumor cell subpopulations as compared to 2D-CRC-PDX cultures.** Consistent with their growth rate, the percentage of proliferative cells in stage (A) II 3D-eCRC-PDX tissues was higher than those in (B) stage III-B and (C) stage IV 3D-eCRC-PDX tissues. Both (D, G) stage II and (E, H) stage III-B 3D-eCRC-PDX tissues maintained both human $CK20^+$ and human $CK20^-$ cell subpopulations over long-term culture; the (I) stage IV 3D-eCRC-PDX tissues could maintain human $CK20^+$ human cell subpopulation over 15 days. In sharp contrast, (D-I) in 2D-CRC-PDX cells, a significant decrease in these cell subpopulations was found after 8 days of culture with the percentages remaining low thereafter. Bars are mean ± SD. Means that do not share a letter are significantly different ($p \leq 0.05$, n = 3 separate batches of cell culture).

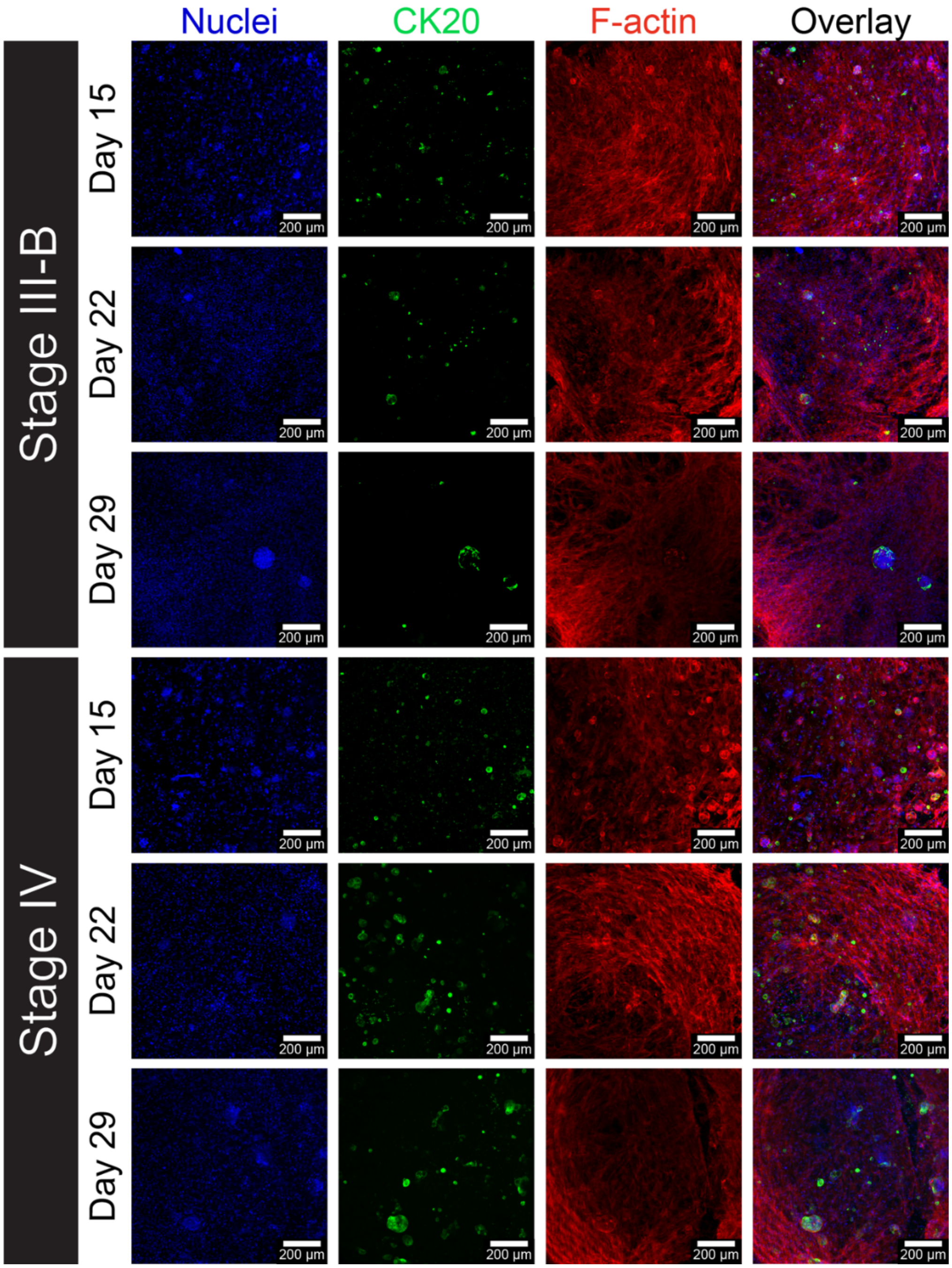


**Supplementary Figure 10. Cellular Staining in 3D-eCRC-PDX Tissues.** Cells were stained with Hoechst 33342 (blue), CK20 (green), and phalloidin (red). $CK20^+$ cells formed colonies, while $CK20^-$ mouse cells exhibited elongation. The density of elongated cells increased from Day 1 to Day 29, indicating a dynamic shift in cellular morphology over time.

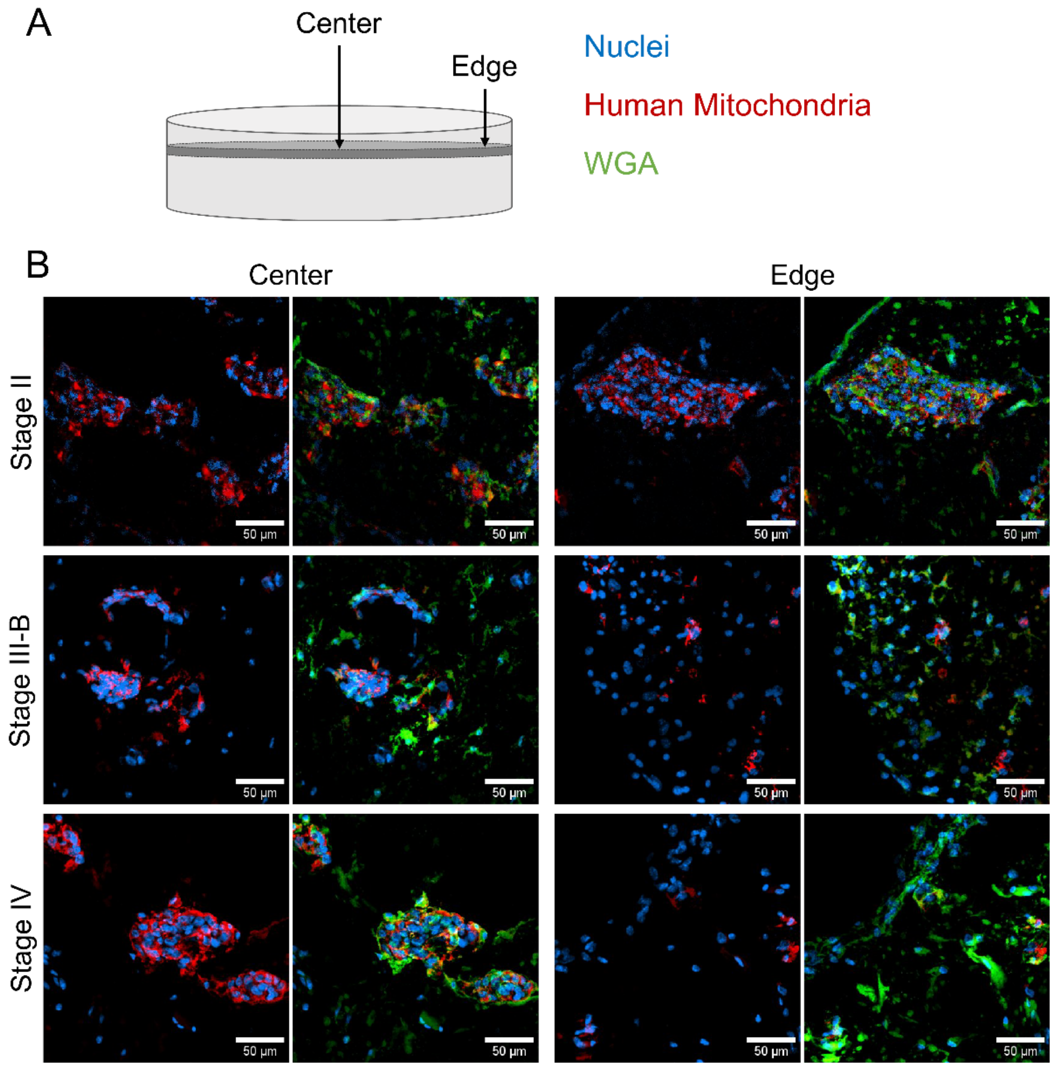


**Supplemental Figure 11**. Immunostaining of sectioned 3D-eCRC-PDX tissues illustrates species-dependent cell populations in each PDX line; similar to flow cytometry analyses, human cells constituted a substantial proportion of the total 3D-eCRC-PDX cell population. A) 3D-eCRC-PDX tissue schematic indicating geometric locations of analysis and a color legend for each fluorescent label employed. B) Confocal microscopy images of immunostained 3D-eCRC-PDX tissue sections (blue indicates cell nuclei, red indicates human mitochondria, and green indicates WGA; scale bars are 50 μm).

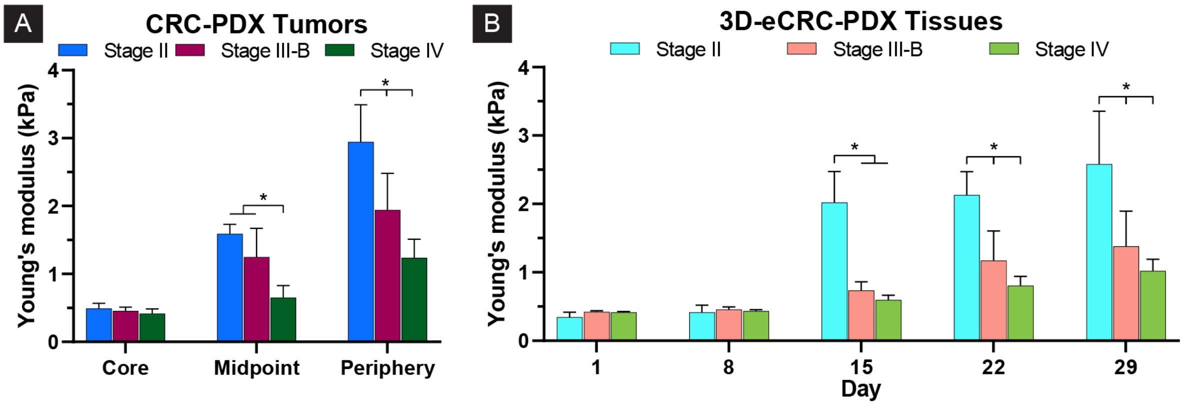


**Supplementary Figure 12. The 3D-eCRC-PDX tissues mimicked the stage-dependent mechanical stiffness of CRC-PDX tumors.** The average stiffness of all batches (pooled data of 3 separate batches of culture) was CRC stage-dependent. (A) The periphery of stage II CRC-PDX tumors had the highest stiffness, followed by those of stage III-B, and then stage IV. (B) Remarkably, the average stiffness of all batches of 3D-eCRC-PDX tissues mimicked the stage-dependent average stiffness of the CRC-PDX tumors. Bars are mean ± SD ($p \leq 0.05$, n = a minimum of 6 tissues from at least 2 separate batches of cell culture).

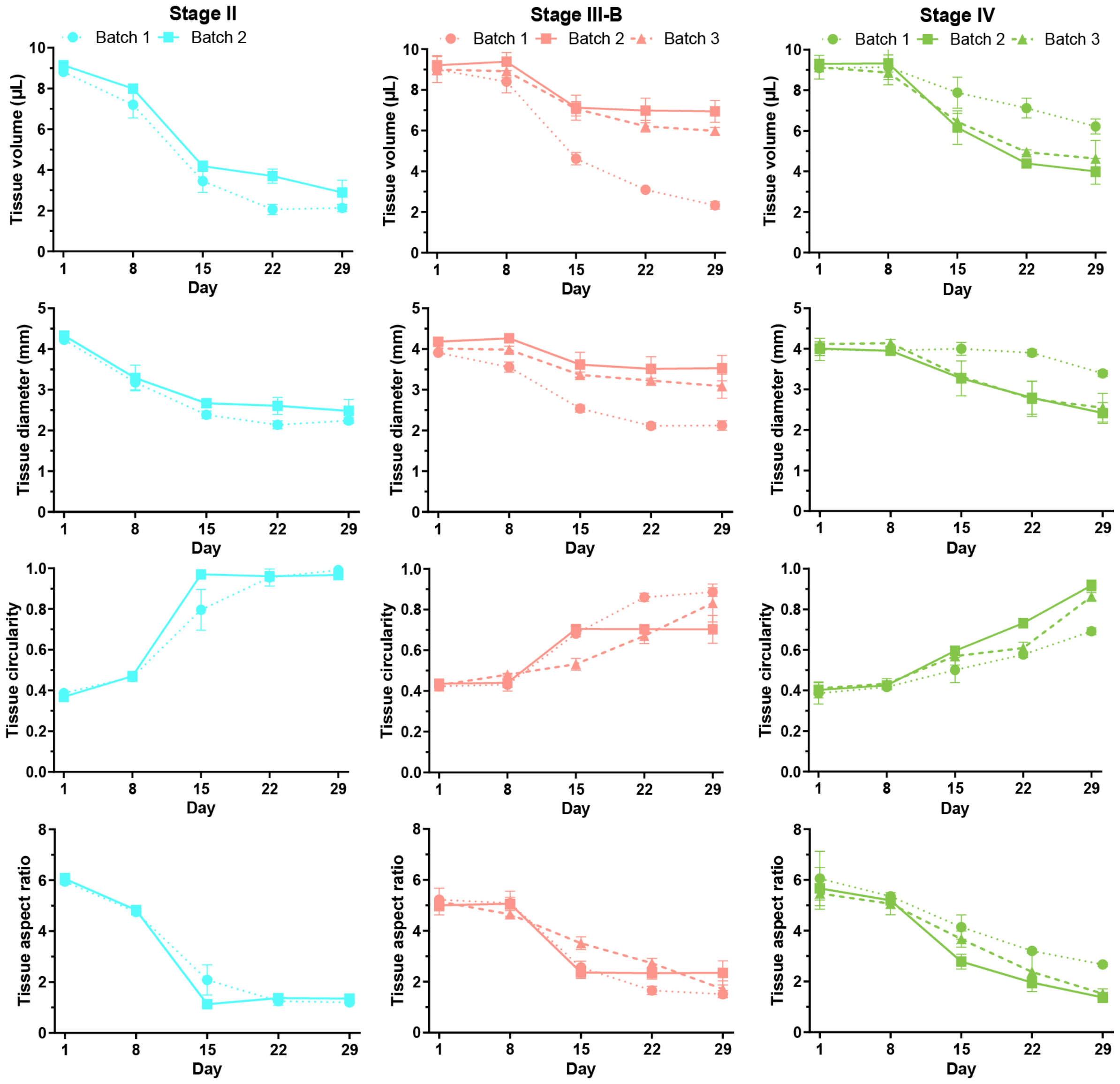


**Supplementary Figure 13. Quantification of size and shape of the 3D-eCRC-PDX tissues for a minimum of two separate batches of cultures.** For each CRC stage, area, diameter, circularity, and aspect ratio of the 3D-eCRC-PDX tissues were quantified over 29 days of culture. For all CRC stages, the volume, diameter, and aspect ratio (side view) decreased and circularity (side view) increased significantly over time; however, the rates of change were different among CRC stages. Data are mean ± SD (n = 3 tissues per time point per batch).

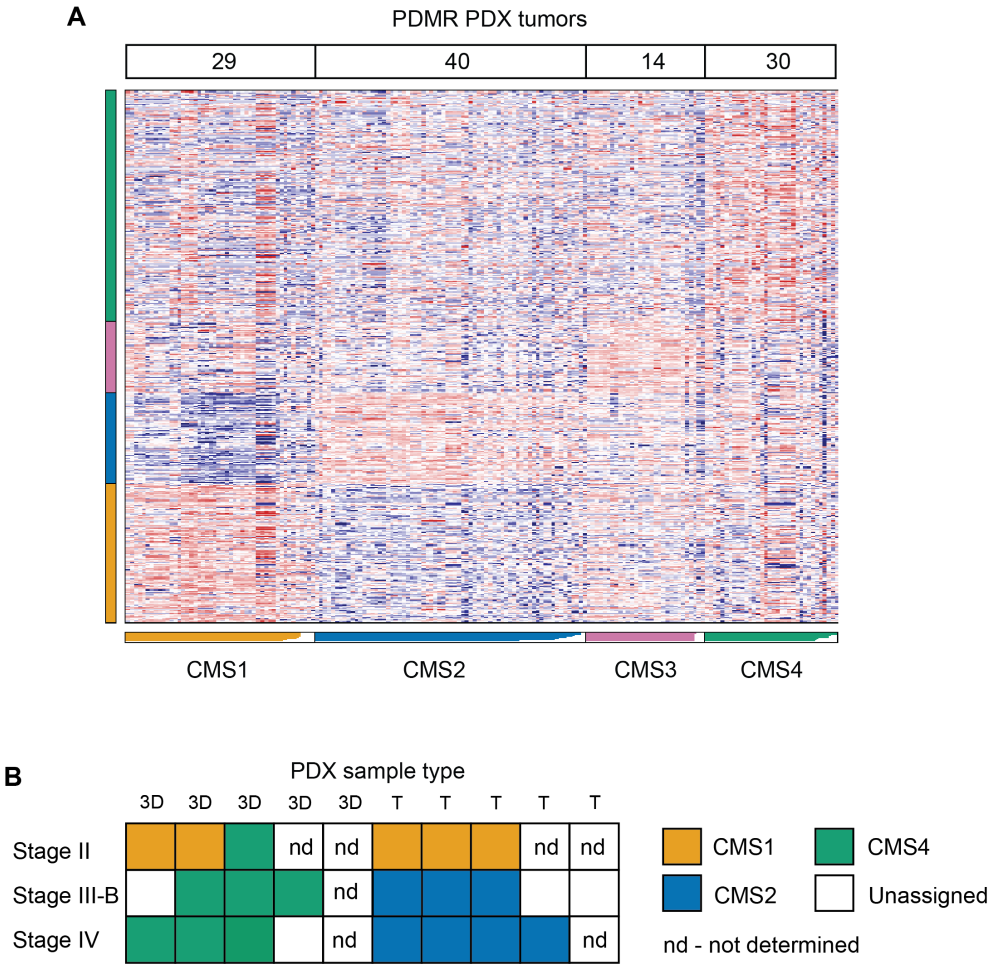


**Supplementary Figure 14. Transcriptomic analysis of 3D-eCRC-PDX tissues demonstrates that 3D-eCRC-PDX tissues Consensus Molecular Subtype is PDX line-dependent.**

(A) Heatmap of CMS classification of 113 colon adenocarcinoma PDX tumors from a cohort of 157 colon adenocarcinoma PDX tumors in the NCI Patient-Derived Models Repository (PDMR) classified as CMS1-4. (B) Diagram of CMS classification. The classification was performed using RNA seq counts data from the PDMR together with 3D-eCRC-PDX tissues (3D) and originating PDX tumors (T) RNA seq counts data.

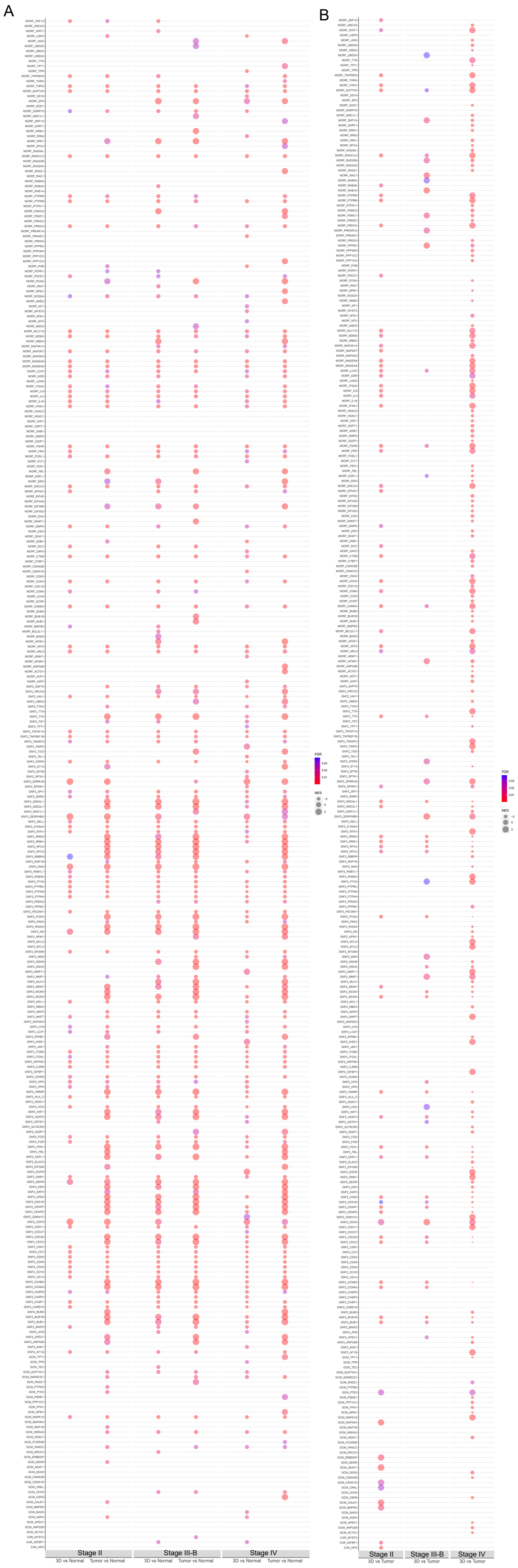
A
B
Stage II
Stage III-B
Stage IV
3D vs Normal
Tumor vs Normal
3D vs Tumor
FDR
NES

**Supplemental Figure 15. GSEA analysis between 3D-eCRC-PDX tissues and CRC-PDX tumors for human and mouse for each stage.** GSEA analysis showing the enriched for cancer-oriented computational genes in (A) 3D-eCRC-PDX tissues and CRC-PDX tumors compared to normal colon tissues (n = 41) and (B) 3D-eCRC-PDX tissues compared to CRC-PDX tumors for each stage. The following three subcollections defined by mining large collections of cancer-oriented expression data are included: Curated Cancer Cell Atlas (3CA), Cancer gene neighborhoods (CGN), and Cancer modules (CM). The bubble size indicates the normalized enrichment score (NES). The bubble color indicates the false discovery rate (FDR).

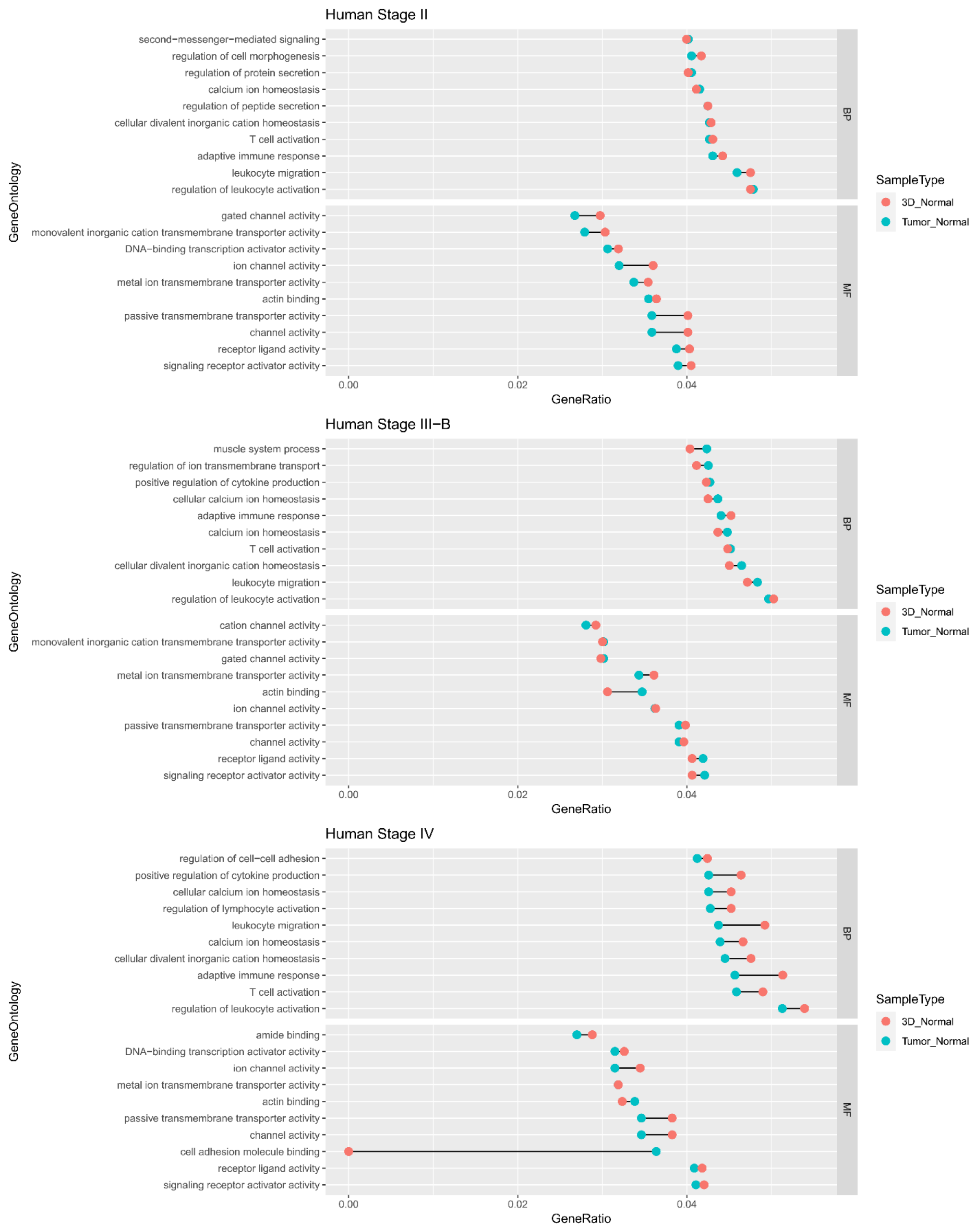


**Supplemental Figure 16. Transcriptomic analysis demonstrated that 3D-eCRC-PDX tissues recapitulated key molecular characteristics of CRC-PDX tumors in a patient line-dependent**

**manner.** Gene ontology analysis of biological process (BP) and molecular functions (MF) enriched in DEGs between CRC-PDX tumors and 3D-eCRC-PDX tissues and normal colon tissues (n = 41) for each stage.

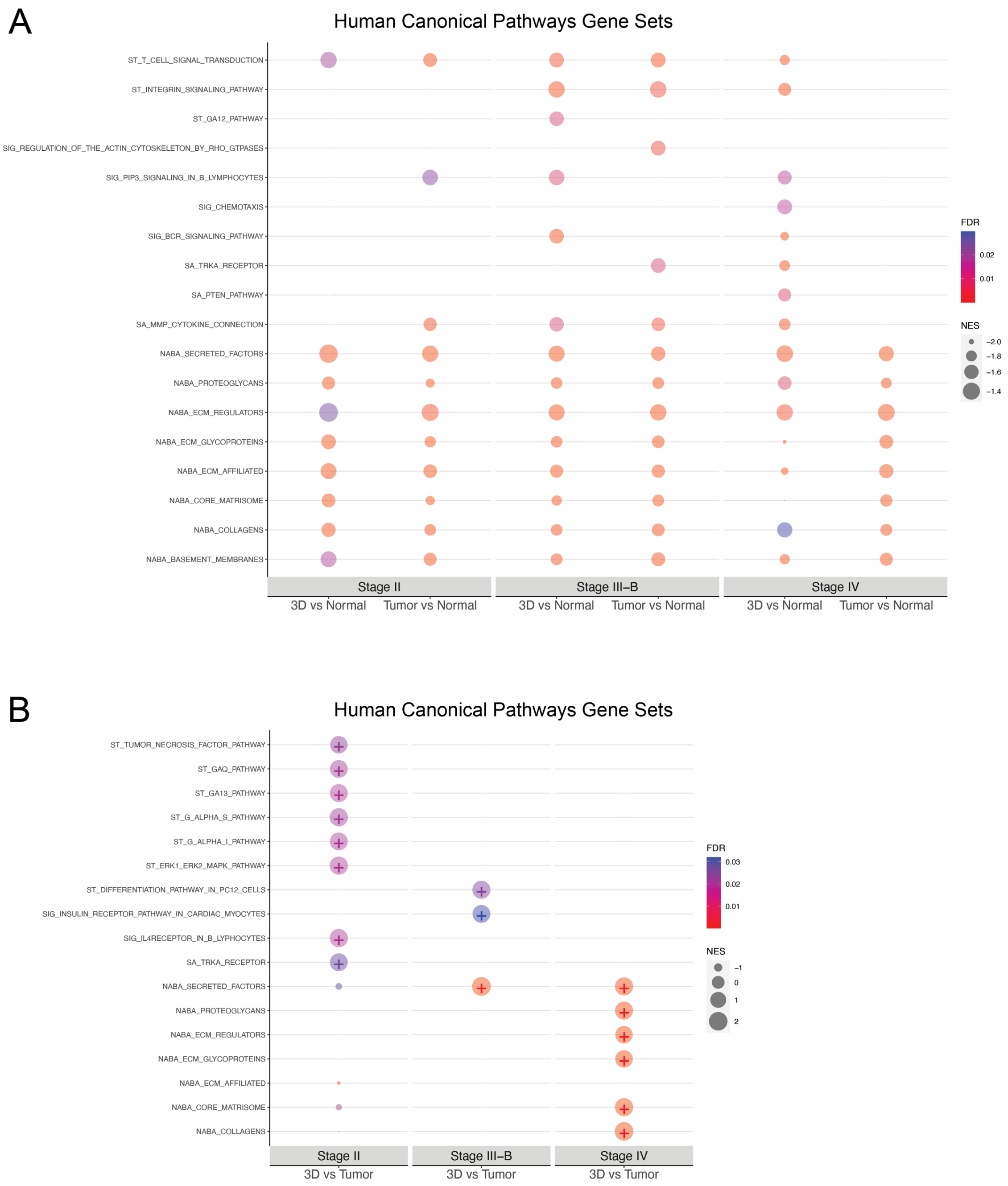


**Supplementary Figure 17. Transcriptomic analysis of 3D-eCRC-PDX tissues and CRC-PDX tumors.** (A) GSEA analysis shows negative enrichment in all 3D-eCRC-PDX tissues and CRC-PDX tumors compared to normal colon tissues (n = 41) for all matrisome gene sets. (B) GSEA analysis shows stage-specific differences of enriched matrisome gene sets in 3D-eCRC-PDX tissues compared to CRC-PDX tumors. The bubble size indicates the normalized enrichment score (NES), and positive enrichment is labeled with a "+" within the bubble. The bubble color indicates the false discovery rate (FDR).

**Supplementary Table 1. Summary of the sequencing reads[Δ] alignment to individual mouse and human reference genomes* for three CRC-PDX lines**

| | Stage II | | Stage III-B | | Stage IV | |
|---|---|---|---|---|---|---|
| | (n=3) | (n=3) | (n=4) | (n=6) | (n=4) | (n=5) |
| | **3D[†]** | **CRC-PDX[‡]** | **3D[†]** | **CRC-PDX[‡]** | **3D[†]** | **CRC-PDX[‡]** |
| **All reads** | 416,126,287 | 503,165,988 | 699,043,154 | 999,265,798 | 647,281,805 | 618,844,880 |
| **Human mapped reads** | 327,161,183 (78.6%) | 452,624,548 (89.6%) | 224,455,276 (32.1%) | 935,849,910 (93.6%) | 110,241,628 (17.0) | 585,358,438 (94.6%) |
| **Mouse mapped reads** | 88,965,104 (21.4%) | 50,541,440 (10.0%) | 474,587,878 (67.9%) | 63,415,888 (6.3%) | 537,040,177 (83.0%) | 33,486,442 (5.4%) |

[Δ]Analysis performed using samtools idxstats tool
*GRCh38 and GRCm38 (mm10)
[†]3D-eCRC-PDX tissues
[‡]CRC-PDX tumors
